# Multimodal PET Defines a ‘Goldilocks’ Thermal Window for Focused Ultrasound Ablation and Immunotherapy Combinations

**DOI:** 10.64898/2026.06.10.731490

**Authors:** Zehra E. F. Demir, Thomas Sherlock, Matthew R. DeWitt, Pouria Talebibarmi, Christian Palacios-Gomez, Alexander L. Kilbanov, Kiel D. Neumann, Shayn M. Peirce, Matthew J. Lazzara, Jonathan R. Lindner, Jiang He, Bijoy Kundu, Natasha D. Sheybani

## Abstract

**Background:** Thermally ablative focused ultrasound (T-FUS) offers a noninvasive, spatially precise strategy for local tumor destruction, with the added potential to remodel tumor architecture and immune dynamics in ways that influence downstream therapeutic delivery and efficacy. Despite promising preclinical and clinical findings, the T-FUS parameters that best balance tumor debulking with preservation of local biologic, e.g. immunotherapy, penetrance remain unclear. Thermal dose, defined by the relationship between tissue heating, exposure duration, and biological effect, is likely a critical determinant of this balance. Excessive thermal dose may eliminate the vascular and stromal features needed to support immunotherapy access, whereas insufficient thermal dose may fail to achieve meaningful cytoreduction. Here, we deploy multimodal PET, contrast-enhanced ultrasound, and tissue profiling to define a “Goldilocks Zone” for T-FUS that balances bulk tumor destruction with immunotherapy delivery.

**Method:** Subtotal T-FUS was applied to 4T1 tumors using three thermal dose regimens resolved by *in silico* modeling. Ablation was quantified by H&E and TTC staining. Post-ablative perfusion and microvascular coverage were assessed by contrast-enhanced ultrasound and immunofluorescence, respectively. Tumor oxygenation was measured by intravenous hypoxyprobe labeling. After T-FUS, mice underwent dynamic [^18^F]-FDG PET and immunoPET with a model tumor-targeted antibody, [^89^Zr]-αCD47, to relate cytoreduction to antibody penetrance. ImmunoPET findings were further evaluated by *ex vivo* biodistribution analysis.

**Results:** *In silico* modeling established three T-FUS regimens that generated distinct thermal dose profiles and were deployed *in vivo* in a solid breast tumor model. Histopathology, perfusion imaging, and hypoxia analysis revealed dose-dependent and dose-divergent biological effects that informed a candidate Goldilocks thermal window. Low thermal dose produced measurable but limited tumor debulking, whereas high thermal dose caused disproportionate functional perfusion collapse. An intermediate thermal dose achieved robust partial ablation, broad hypoxia relief, and preservation of residual tumor physiology sufficient to support antibody access. Dynamic [^18^F]-FDG PET confirmed a marked reduction in metabolically active tumor burden after Goldilocks T-FUS. Serial [^89^Zr]-αCD47 immunoPET showed that bulk antibody signal was maintained after ablation, and integration of immunoPET with matched [^18^F]-FDG PET revealed approximately 3-fold enrichment of antibody exposure within the residual viable tumor compartment of ablated tumors. These findings demonstrate that appropriately tuned thermal ablation can debulk tumor while preserving, and potentially concentrating, immunotherapy access within the remaining targetable tumor niche.

**Conclusion:** This study identifies thermal dose as a critical consideration for T-FUS immunotherapy combinations and establishes a PET-informed framework for balancing cytoreduction with therapeutic delivery. Rather than functioning solely as a local debulking modality, we demonstrate that T-FUS can be tuned to yield a post-ablation tumor state that remains accessible to large biologics. These findings provide timely, translationally relevant guidance for tailoring T-FUS regimens to achieve local tumor destruction while preserving an immunotherapy-permissive niche for combination treatment.

## INTRODUCTION

Locally ablative techniques, such as laser ablation, cryoablation, radio-frequency ablation, irreversible electroporation, and microwave ablation, are increasingly being explored as alternatives or complements to conventional cancer treatment^1–3^. These approaches may be especially valuable for patients who are poor surgical candidates due to age, comorbidities, or tumor location. However, many of these modalities remain invasive and carry important procedural limitations. Thermally ablative focused ultrasound (T-FUS), by contrast, offers a truly noninvasive, non-ionizing, and spatially precise strategy for local tumor destruction - with the added potential to modulate tumor architecture and the immune response in ways that influence downstream therapeutic delivery and efficacy^4^. With its completely incisionless approach and safe treatment profile, T-FUS induces localized cytoreduction and heat-induced stress signaling through controlled deposition of acoustic energy. By rapidly raising tissue temperature within a confident focal volume, T-FUS yields instantaneous coagulative necrosis while neighboring periablative margins undergo sub-ablative hyperthermia within a volume defined by parameters such as frequency, power, and duration of sonications. Decades of technical development have underscore the exponential clinical adoption of T-FUS across a range of oncological indications^4,5^, with numerous commercial devices available.

In addition to tumor debulking, T-FUS has demonstrated immunomodulatory benefits^6–8^. To this end, it has emerged as a burgeoning strategy for potentiate immunotherapies such as checkpoint inhibitors^9–12^, co-stimulatory immunotherapy^12^, and immune adjuvants^10,11,13,14^ in both preclinical and clinical studies^4,10,11^. Notably, several completed (e.g. NCT06258330, NCT03237572, NCT04116320) and ongoing (e.g. NCT06964906, NCT06472661, NCT07126093) clinical trials are investigating drug combinations with T-FUS, chief among which are biologics such as immune-directed antibodies.

Despite promising preclinical and clinical evidence to date, whether T-FUS parameters can be tuned to balance generous tumor cytoreduction with maintained local antibody penetrance remains undefined. Limited ablative immuno-modulation studies have evaluated the impact of scanning density^15^ or ablation type^11,16,17^. Meanwhile, numerous T-FUS immuno-oncology clinical trials (NCT04116320, NCT03237572, NCT04796220, NCT06472661) have converged on the practice of subtotal ablation. Even though patients understandably may favor the intuitive appeal of complete tumor eradication, this underscores the collective postulation of a “Goldilocks Zone” for thermal ablation that balances debulking and cooperation with tumor-directed immunotherapy (Figure 1A). To interrogate this zone, we focus on the understudied principle of thermal dose as a critical determinant of biological outcomes. We first disentangle the influence of thermal dose on features of tumor architecture and then proceed to examine how a thermal dose within the postulated “Goldilocks Zone” impacts penetrance of a model tumor-targeted immunotherapy, mCD47, using a multimodal positron emission tomography (PET) approach. These findings of this study offer provocative, translationally relevant insights regarding the interplay between thermal destruction and macromolecular therapeutic penetrance antibody penetrance in solid cancers.

**Figure 1:**
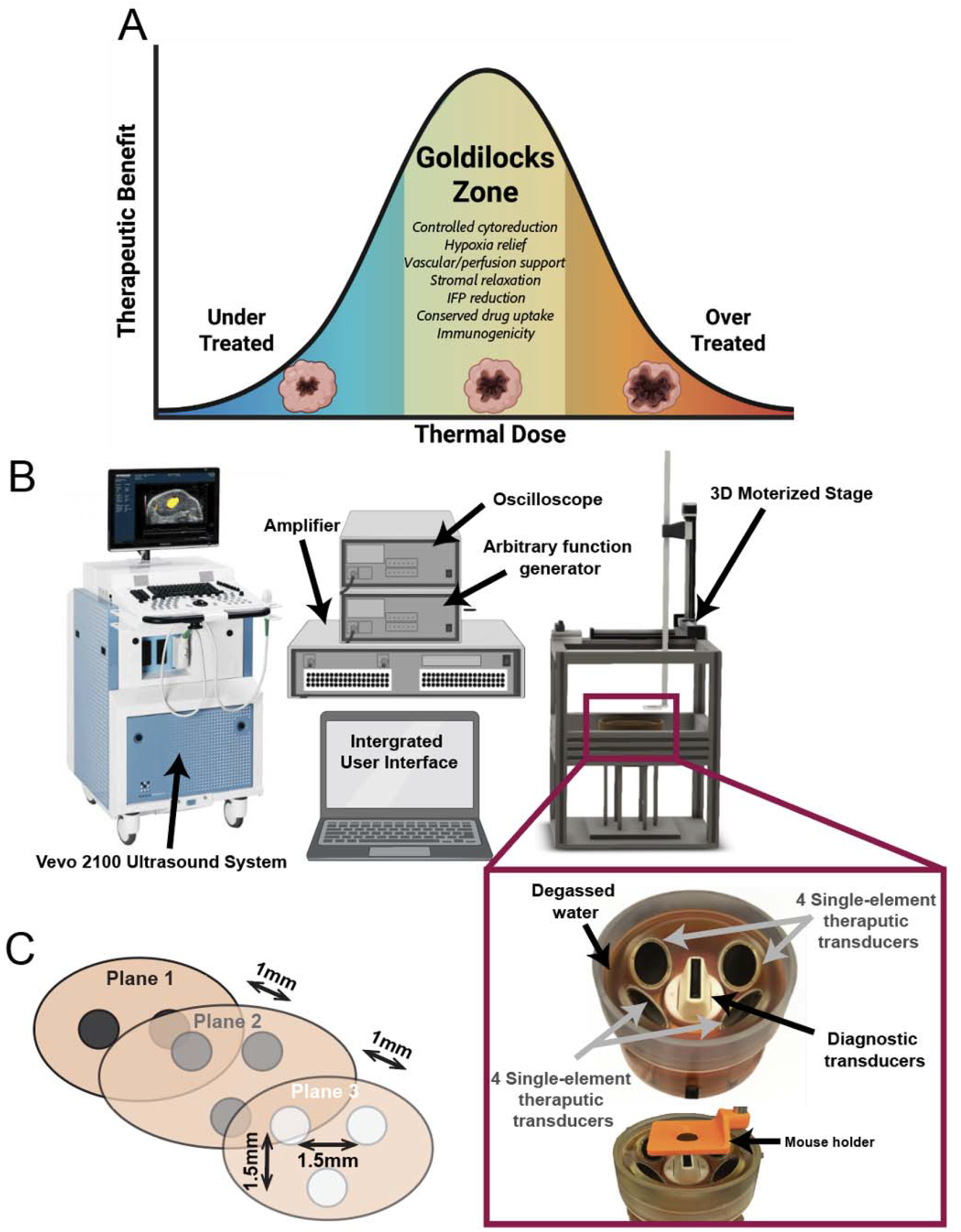
Conceptual and experimental framework for defining “Goldilocks Zone” for FUS thermal ablation. A) Proposed concept of “Goldilocks Zone” in which T-FUS achieves controlled tumor cytoreduction while preserving residual tumor and stromal features that may support therapeutic delivery and anti-tumor immune activity. Undertreatment may provide insufficient tumor debulking, whereas overtreatment may compromise the vascular and stromal physiology needed for therapeutic access. B) Illustration of custom ultrasound-guided FUS system. C) Example schematic of multi-plane T-FUS sonication pattern.

## RESULTS

### In silico simulation establishes three distinct thermal dose profiles for partial T-FUS yielding dose-dependent increases in maximum temperature and ablation volume

A 3D computational model was developed to simulate three discrete partial T-FUS treatment scenarios corresponding to low, targeted-moderate, and high/near-complete thermal doses (T-FUS_Low_, T-FUS_Mid_, T-FUS_High_). The geometry consisted of layered domains representing skin and underlying tumor tissue (Table S1), with dimensions chosen to approximate the experimental custom ultrasound-guided FUS system configuration (Figure 1B, Figure S1A). Furthermore, in all cases, heating was localized around the acoustic focus, with the maximum temperature increasing proportionally with input amplitude (Figure 2A-C). The peak focal temperature achieved in the tumor over T-FUS sonication period increase in a dose-dependent manner, with the area under the curve from 0-15 seconds (AUC_0-15s_) recapitulating this finding (Figure 2D-E). To further explore sonication duration, additional simulations were performed for the medium input amplitude (T-FUS_Mid_) with exposure times of 5s and 25s. The temperature fields and ablation volumes increased as duration of sonication increased (Figure S2). In all, these *in silico* results established and validated three distinct thermal doses for ablation that were carried forward in murine studies.

**Figure 2:**
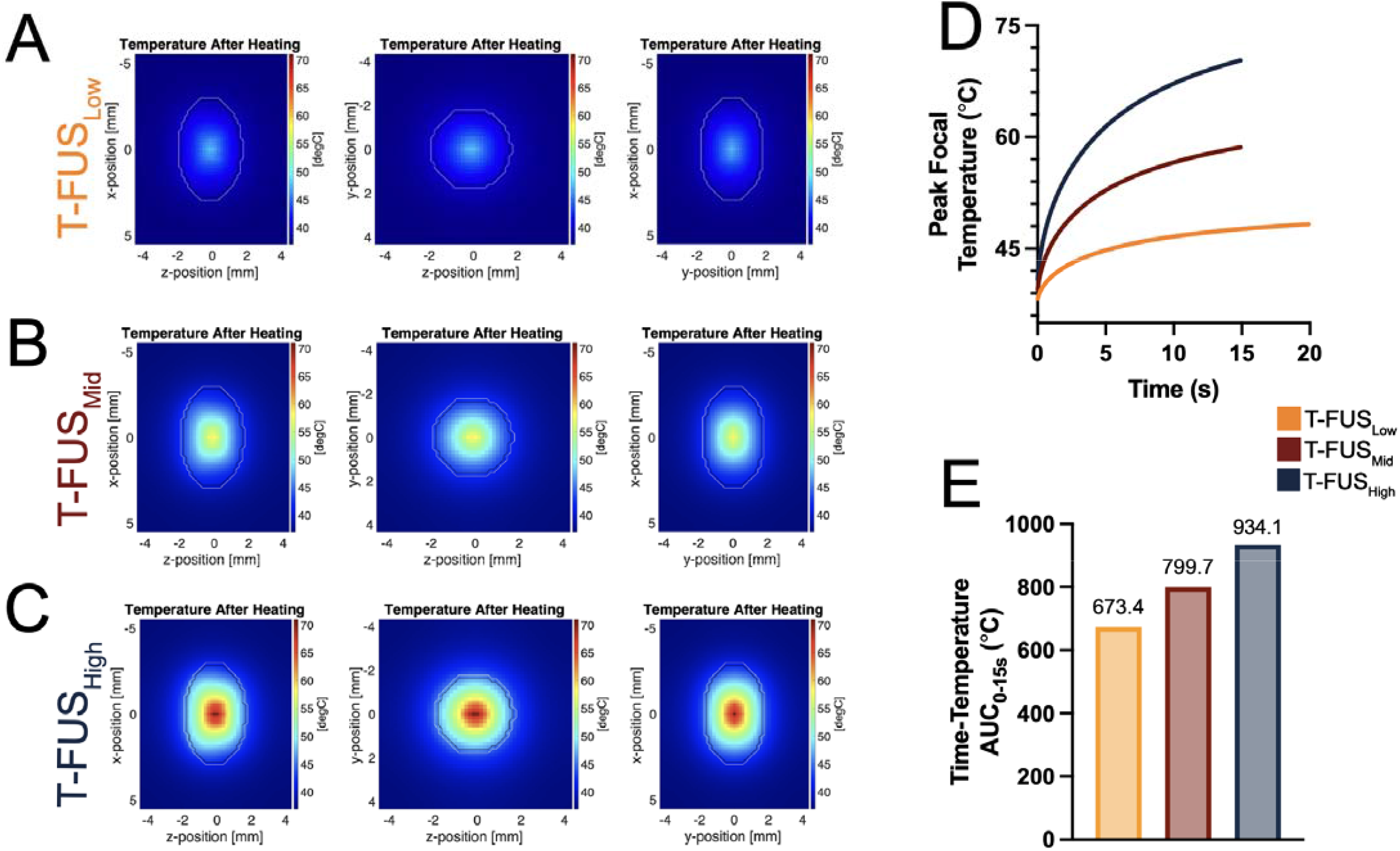
Computational modeling establishes dose-escalated T-FUS regimens with graded focal thermal exposures. A-C) *In silico* temperature maps for T-FUS_Low_, T-FUS_Mid_, and T-FUS_High_ immediately following sonication, shown in the xz, yz, and xy planes through the tumor center. D) Predicted peak focal temperature achieved in tumor for each T-FUS regimen. E) Area under the peak temperature-time curve from 0-15 seconds (AUC_0-15s_), demonstrating increasing thermal exposure across T-FUS_Low_, T-FUS_Mid_, and T-FUS_High_ conditions.

### T-FUS thermal doses yield graded tumor ablation and uniformly relieve hypoxia within the local tumor microenvironment

After establishing three distinct thermal dose regimens via *in silico modeling*, we next sought to evaluate these regimens *in vivo* using the 4T1 triple negative mammary carcinoma model. To assess the degree of ablation, tumors were stained at 8 hours after T-FUS treatment using 2,3,5-triphenyltetrazolium chloride (TTC) stain and 3 days post-treatment with hematoxylin and eosin (H&E) (Figure 3A). Qualitatively, TTC staining revealed graded reduction of metabolically active regions with increasing thermal dose (Figure 3B). H&E staining further revealed the details of coagulative necrosis in the ablative zone of T-FUS treated tumors and a surrounding transition zone of heat-mediated damage. By comparison, controls displayed baseline necrosis of entirely distinct patterning and degree (Figure 3C).

**Figure 3:**
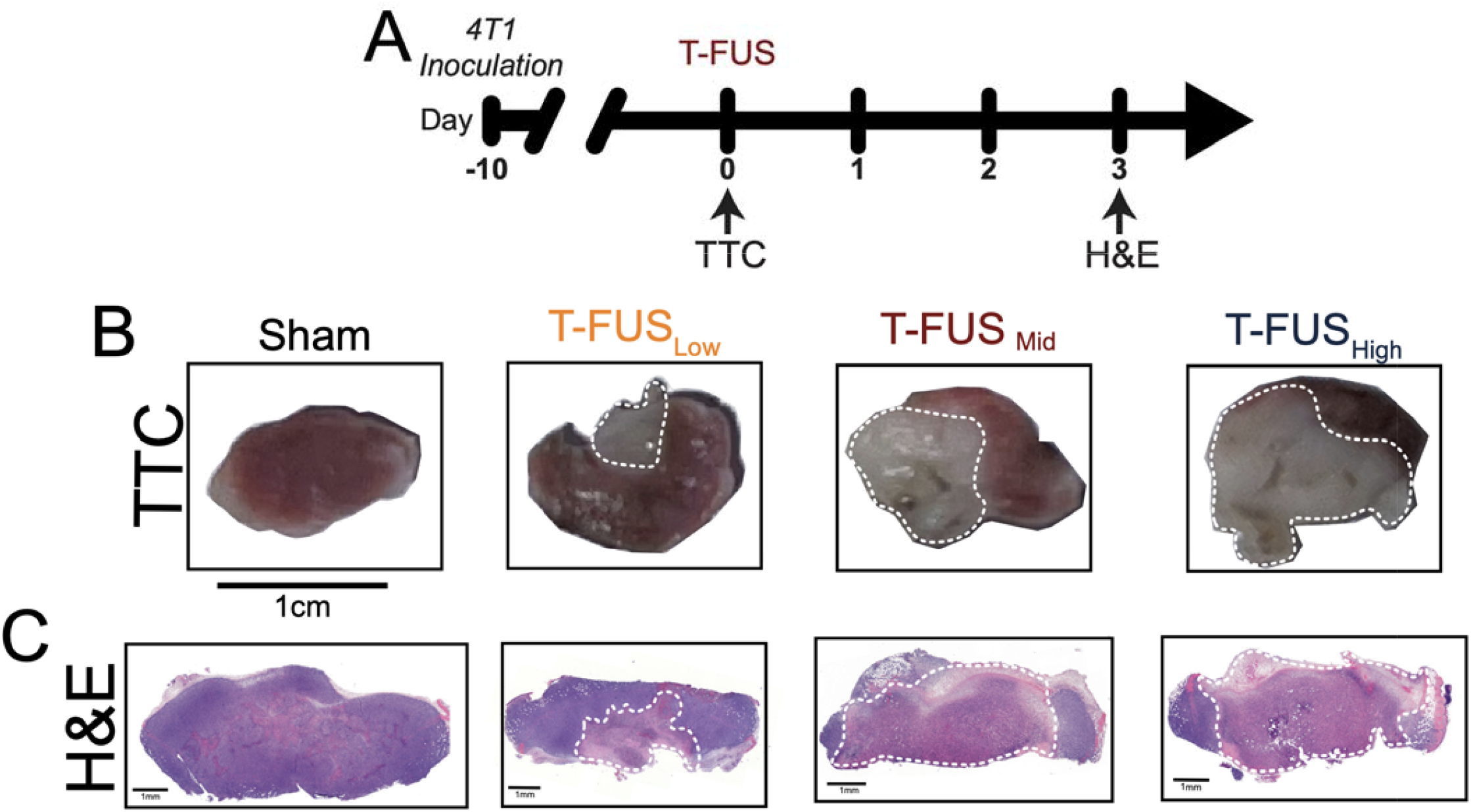
T-FUS thermal doses yield distinct degrees of partial tumor debulking via metabolic staining and histology. A) Experimental timeline for T-FUS treatment and subsequent *ex vivo* tissue analysis. B) Representative triphenyl tetrazolium chloride (TTC)-stained tumors 24 hours after sham or T-FUS treatment. Arrows indicate non-metabolically active regions consistent with graded thermal ablation. C) Representative hematoxylin and eosin (H&E) stained tumor sections 3 days after sham or T-FUS treatment. Arrows indicate regions of thermal injury and necrosis, which increase across dose-escalated T-FUS regimens.

To investigate the degree and spatial patterning of hypoxia following thermal ablation *in vivo*, we administered intraperitoneal (i.p.) pimonidazole (Hypoxyprobe) at a terminal time point for assessment by immunofluorescence (Figure 4A). Sham controls exhibited abundant and spatially distributed hypoxyprobe staining. This signal was comparatively abolished, and where present, confined to patches localized within tumor margins, suggesting that T-FUS acutely and dramatically reduces hypoxic burden (Figure 4B).

**Figure 4:**
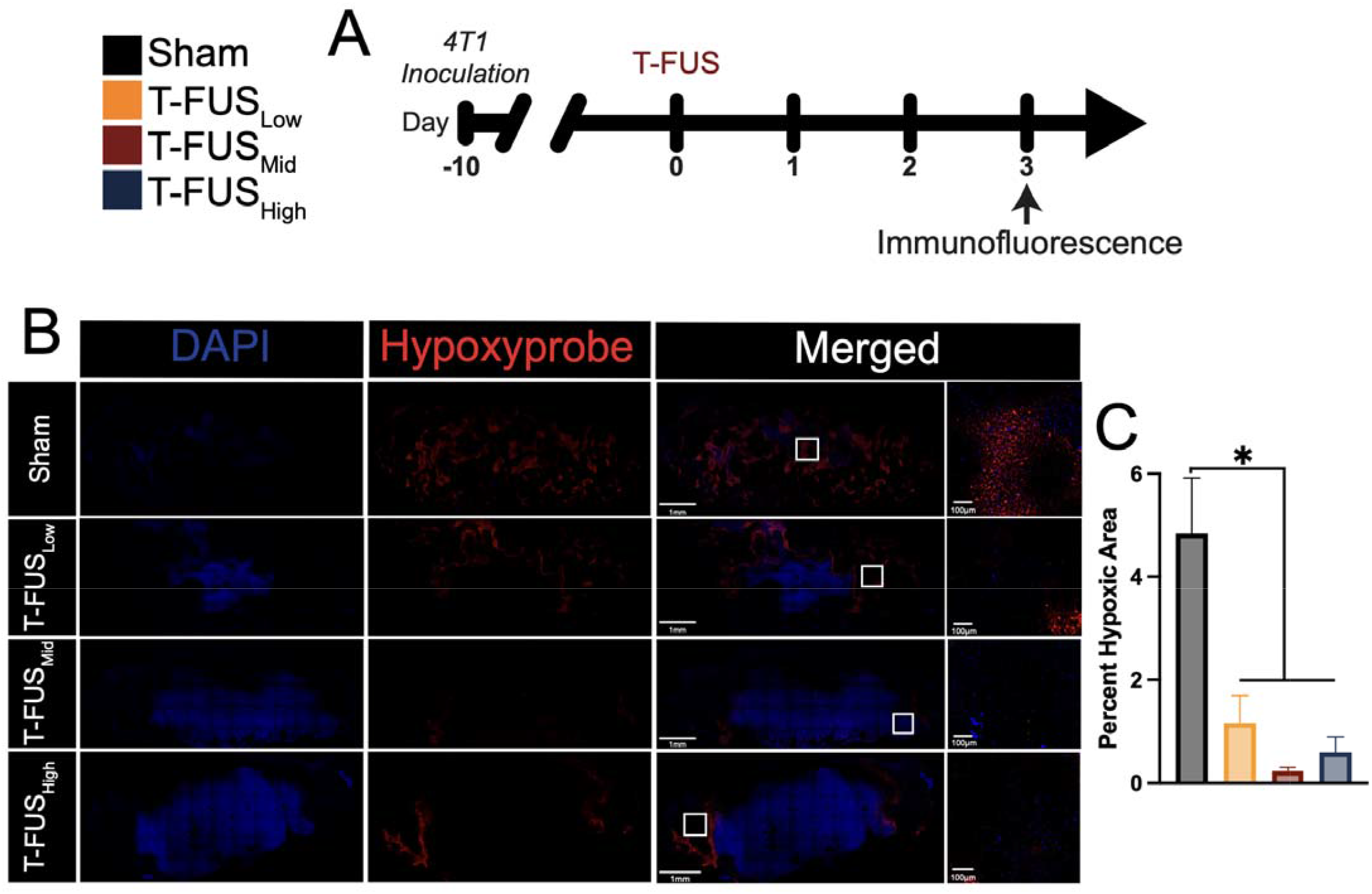
T-FUS reduces tumor hypoxia across thermal dose regimens without a strictly dose-dependent response. A) Experimental timeline for T-FUS treatment and immunofluorescence analysis. B) Representative immunofluorescence images of nuclei (DAPI, blue) and hypoxic regions (hypoxyprobe, red) in sham and T-FUS-treated tumors across low, mid and high thermal dose regimens. Insets show higher-magnification views of the indicated regions (white box). C) Quantification of percent hypoxic area. *p<0.05 vs indicated groups: Sham vs. T-FUS_Low_ (p = 0.0006), Sham vs T-FUS_Mid_ (p < 0.0001), Sham vs T-FUS_High_ (p = 0.0001). Significance was assessed by one-way ANOVA. n=3 per group; with 3 tumor sections per mouse.

Specifically, T-FUS_Low_, T-FUS_Mid_, and T-FUS_High_ tumors exhibited approximately 4.1-, 20.9-, and 8.1-fold decreases in hypoxyprobe staining relative to sham tumors, respectively (Figure 4C). Together, these findings reveal that the biological consequences of T-FUS are not strictly dose-linear. Increasing thermal dose progressively enhanced tumor debulking, yet hypoxia was relieved across T-FUS-treated tumors without a clear monotonic relationship to dose. We therefore next investigated whether tumor vascular density and perfusion represented more dose-sensitive features of the post-ablation microenvironment.

### T-FUS alters tumor microvascular architecture and reduces functional perfusion in a thermal dose-dependent manner

In tandem with hypoxia staining, we probed the tumor endothelium (endomucin) via immunofluorescence staining to determine the relationship between thermal dose and microvascular architecture (Figure 5A). For all T-FUS groups, endomucin staining was reduced, particularly within the ablative zone, compared to the sham group. Qualitatively, a diminution in positive staining was evident from sham to T-FUS conditions (Figure 5B). Quantification of mean vessel area and mean vessel percentage across slices revealed that T-FUS_Low_, T-FUS_Mid_, and T-FUS_High_ conditions reduced average vessel area by approximately 1.5-, 4.4-, and 2.0-fold relative to sham controls, respectively (Figure 5C). Furthermore, T-FUS_Low_, T-FUS_Mid_, and T-FUS_High_ markedly decreased average vessel percentage, by approximately 1.5-, 3.5-, and 1.9-fold relative to sham controls, respectively (Figure 5D).

**Figure 5:**
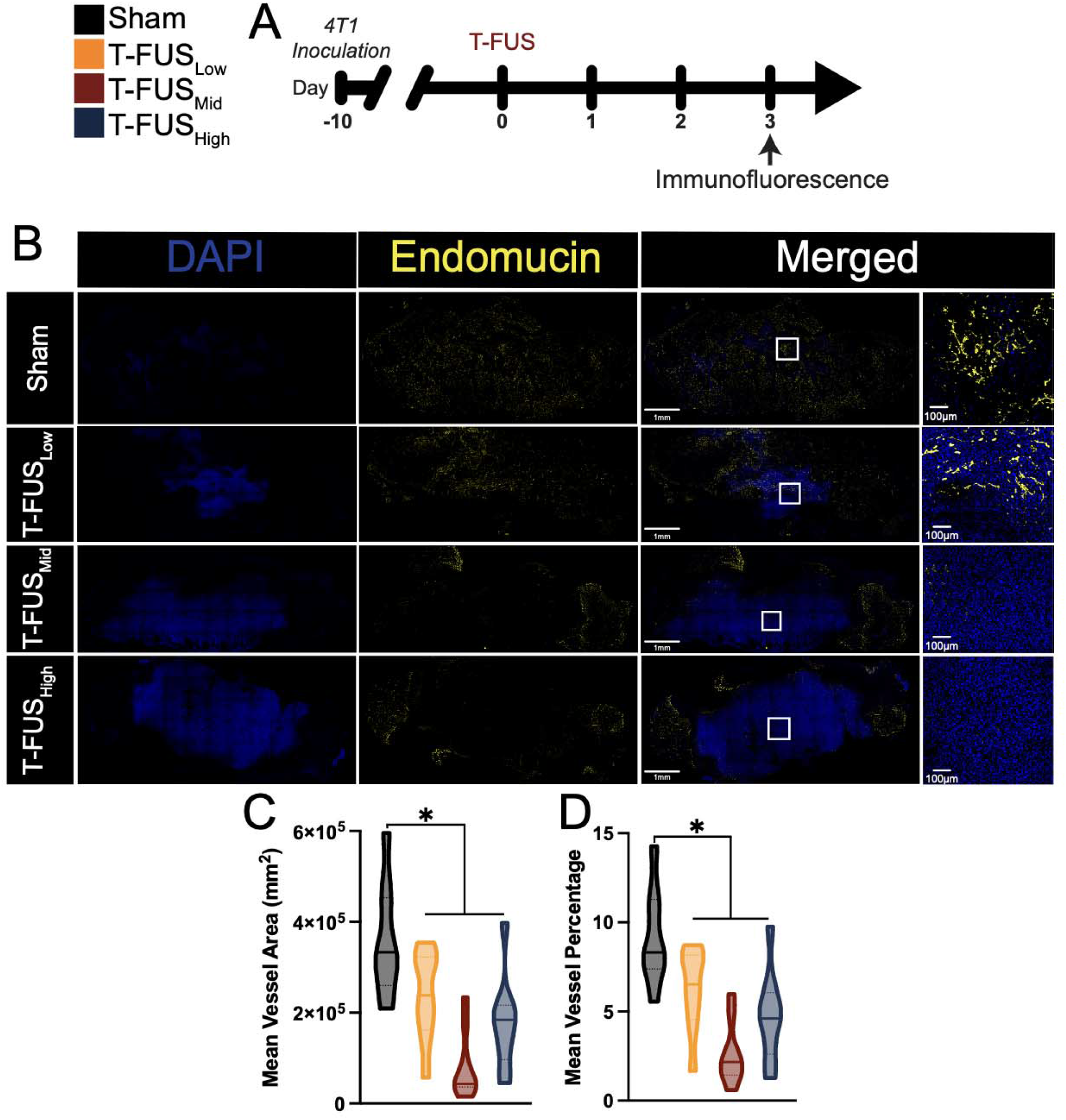
T-FUS disrupts tumor microvascular architecture across thermal dose regimens. A) Experimental timeline for T-FUS treatment and immunofluorescence analysis. B) Representative immunofluorescence images of nuclei (DAPI, blue) and endothelial structures (endomucin, yellow) in sham and T-FUS-treated tumors across low, mid and high thermal dose regimens. Insets show higher-magnification views of the indicated regions (white box). C) Quantification of mean endomucin-positive vessel area. *p<0.05 vs indicated groups: Sham vs. T-FUS_Low_ (p = 0.0472), Sham vs T-FUS_Mid_ (p < 0.0001), Sham vs T-FUS_High_ (p = 0.0019). D) Quantification of mean endomucin-positive vessel percentage. *p<0.05 vs indicated groups: Sham vs. T-FUS_Low_ (p = 0.0385), Sham vs T-FUS_Mid_ (p < 0.0001), Sham vs T-FUS_High_ (p = 0.0013). Significance was assessed by one-way ANOVA. n=3 per group; with 3 tumor sections per mouse.

To determine whether the observed disruption in tumor microvascular architecture translated into functional changes in blood flow, 4T1-bearing mice underwent contrast-enhanced ultrasound (CEUS) perfusion imaging immediately following T-FUS (Figure 6A). Consistent with microscopy findings, we observed that T-FUS uniformly reduces tumor perfusion relative to sham controls (Figure 6B). Quantification of microbubble flux rate and peak intensity from CEUS loops revealed thermal dose-dependent reductions (Figure 6C-D). Specifically, T-FUS_Low_, T-FUS_Mid_, and T-FUS_High_ markedly decreased the rate at which the microbubbles entered the tumor by approximately 2.3-, 4.2-, and 23.6-fold relative to the sham control (Figure 6C). Where as the T-FUS_Low_ condition displayed a trend toward reduced peak intensity, the higher thermal doses (T-FUS_Mid_, and T-FUS_High_) invoked significant reduction in peak intensity compared to shams (Figure 6D); specifically, T-FUS_Low_, T-FUS_Mid_, and T-FUS_High_ exhibited approximately 1.5-, 1.8-, 63.2-fold decreases in the peak intratumoral microbubble signal intensity relative to shams, respectively. Taken together, these findings indicate that T-FUS produces coupled changes in tumor microvasculature and contrast-defined perfusion, with the most pronounced loss of functional perfusion observed after the highest thermal dose.

**Figure 6:**
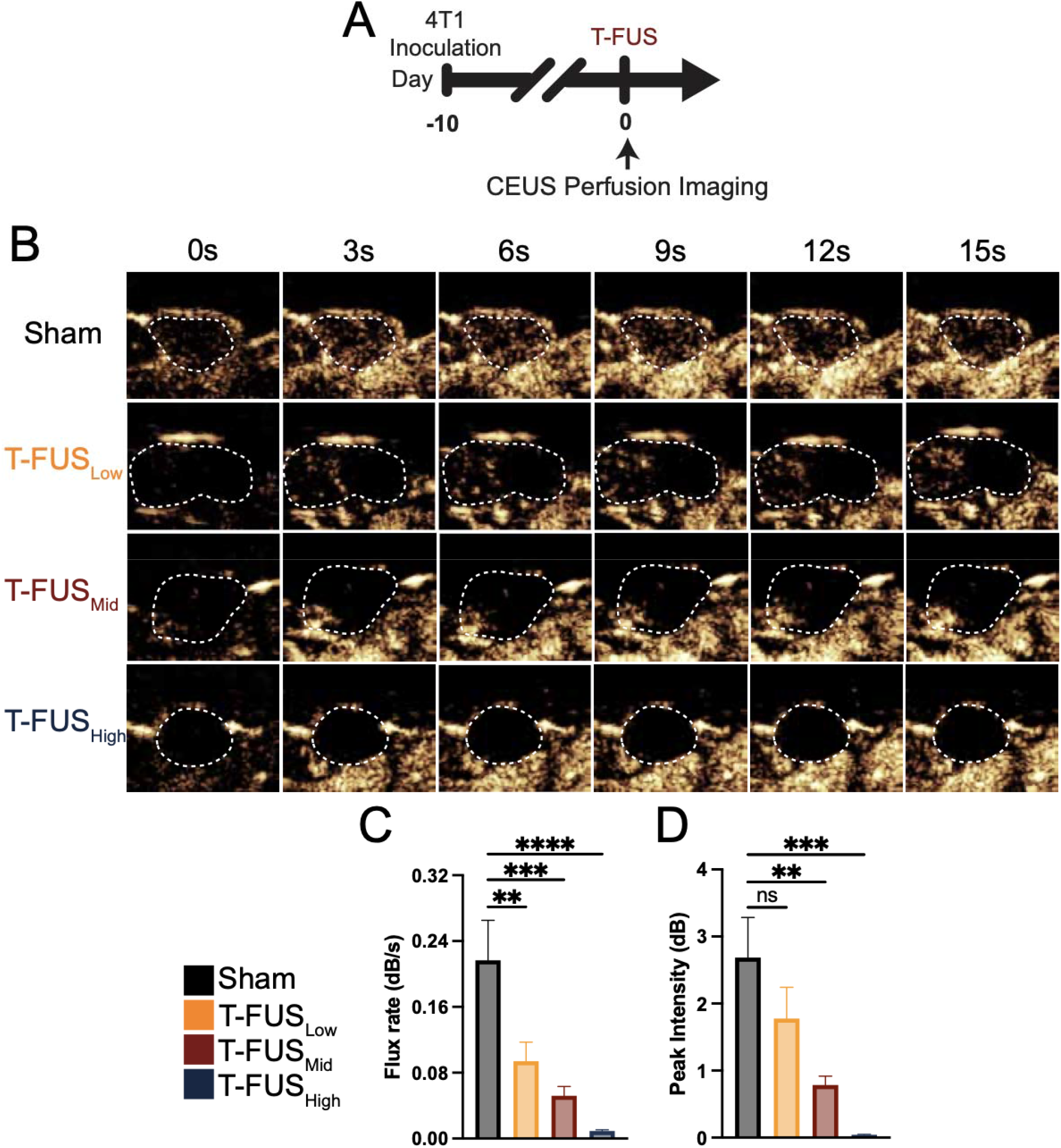
Contrast-enhanced ultrasound perfusion imaging reveals thermal dose-dependent diminution of functional tumor perfusion after T-FUS. A) Experimental timeline for T-FUS treatment and CEUS perfusion imaging performed immediately after treatment. B) Representative CEUS images acquired at serial time points following contrast administration in sham and T-FUS-treated tumors across low, mid and high thermal dose regimens. Dashed white outlines indicate tumor boundaries. C) Quantification of flux rate, reflecting the rate of contrast arrival during the initial upslope phase. Sham vs. T-FUS_Low_ (p = 0.0034), Sham vs T-FUS_Mid_ (p = 0.0002), Sham vs T-FUS_High_ (p < 0.0001). D) Quantification of peak intensity, reflecting maximum intratumoral contrast signal after bolus administration. Sham vs. T-FUS_Low_ (p = 0.1988), Sham vs T-FUS_Mid_ (p = 0.0044), Sham vs T-FUS_High_ (p = 0.0001). Significance was assessed by one-way ANOVA. n=4-6 per group.

These analyses provided the rationale for advancing T-FUS_Mid_ for subsequent PET-based studies. T-FUS_Low_ produced measurable biological effects but did not maximize tumor debulking, whereas T-FUS_High_ caused a disproportionate reduction in functional perfusion, consistent with overtreatment state in which post-thermal tissue changes (e.g. edema) and vascular compromise could limit delivery to residual tumor. In contrast, T-FUS_Mid_ achieved robust partial ablation and hypoxia relief while avoiding the severe perfusion deficit observed at the highest thermal dose. We therefore selected T-FUS_Mid_ as the operational ‘Goldilocks’ condition for testing whether effective tumor cytoreduction can be achieved while preserving sufficient residual tumor physiology to support local immunotherapy delivery.

### [^18^F]-FDG PET confirms volumetric reduction in viable tumor burden following Goldilocks T-FUS

Having identified T-FUS_Mid_ as the operational Goldilocks condition, we next sought to determine whether this regimen produced a measurable reduction in viable tumor burden using a volumetric, non-terminal imaging readout. To this end, 4T1 tumor-bearing mice were administered [^18^F]-fluorodeoxyglucose (FDG) 4 hours after T-FUS_Mid_ treatment and underwent dynamic PET/CT imaging to quantify residual metabolic activity within the tumor (Figure 7A). Although [^18^F]-FDG is not tumor-specific, reduced intratumoral FDG uptake provides a clinically translatable measure of treatment-induced loss of metabolically active tumor tissue following ablation^18–20^. Qualitative assessment of decay-corrected [^18^F]-FDG PET images revealed visibly reduced intratumoral signal in T-FUS_Mid_-treated tumors compared to sham controls (Figure 7B). Quantification of tumor mean standardized uptake values from dynamic PET images further demonstrated a sustained reduction in metabolic activity across the 60-minute acquisition period (Figure 7C). Overall, T-FUS_Mid_ reduced intratumoral [^18^F]-FDG signal by approximately 68% reduction relative to sham controls, consistent with substantial loss of metabolically active tumor tissue following partial thermal ablation. The separation between groups was most pronounced during the final 15 minutes of imaging, during which T-FUS_Mid_-treated tumors exhibited a 73% reduction in metabolic signal than sham tumors (Figure 7D). Together, these PET findings provide volumetric confirmation that T-FUS_Mid_ achieves effective metabolic tumor debulking within the proposed Goldilocks thermal window.

**Figure 7:**
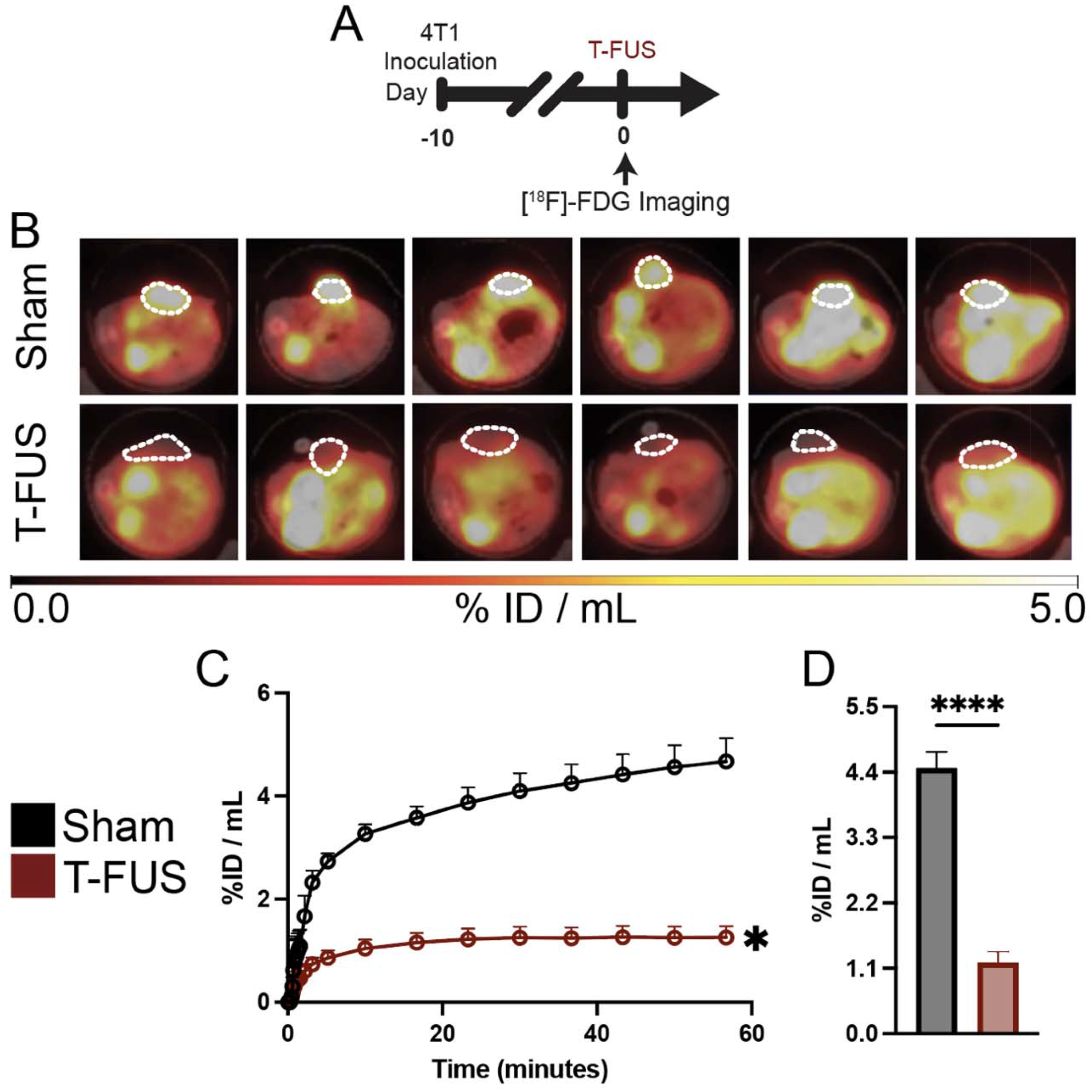
Dynamic [^18^F]-FDG PET imaging confirms volumetric reduction of metabolically active tumor burden following T-FUS_Mid_. A) Experimental timeline for T-FUS treatment and dynamic FDG-PET/CT imaging. B) Representative decay-corrected [^18^F]-FDG PET/CT images showing intratumoral FDG signal in sham and T-FUS tumors (white outline). C) Average tumor time-activity curves over the 60-minute dynamic acquisition, reported as %ID/mL. Significance assessed by two-way ANOVA, followed by Sidak multiple comparison correction. Sham vs T-FUS (p =0.0398). D) Quantification of tumor [^18^F]-FDG signal during final 15 minutes of dynamic imaging. Significance assessed by unpaired t-test. Sham vs T-FUS (p < 0.0001).

### ImmunoPET reveals maintained targeted antibody access following Goldilocks T-FUS

To determine whether T-FUS_Mid_ compromises tumor-targeted antibody delivery, 4T1 tumor-bearing mice received [^89^Zr]-αCD47 one day after treatment and underwent serial PET/CT imaging from day 1 through day 3 post-ablation (Figure 8A). Whole-tumor immunoPET analysis showed that intratumoral [^89^Zr]-αCD47 signal was maintained following T-FUS_Mid_, with no significant differences in longitudinal SUV or AUC compared to sham controls (Figure 8B,E). Linear regression analysis further demonstrated comparable rates of tracer accumulation between groups (Figure 8C), and analogous to the sham condition, [^89^Zr]-αCD47 signal within T-FUS_Mid_-treated tumors increased from day 1 to days 2 and 3 after tracer administration (Figure 8D). These data indicate that partial thermal ablation within the Goldilocks window does not acutely preclude tumor access by a large tumor-targeted antibody.

**Figure 8:**
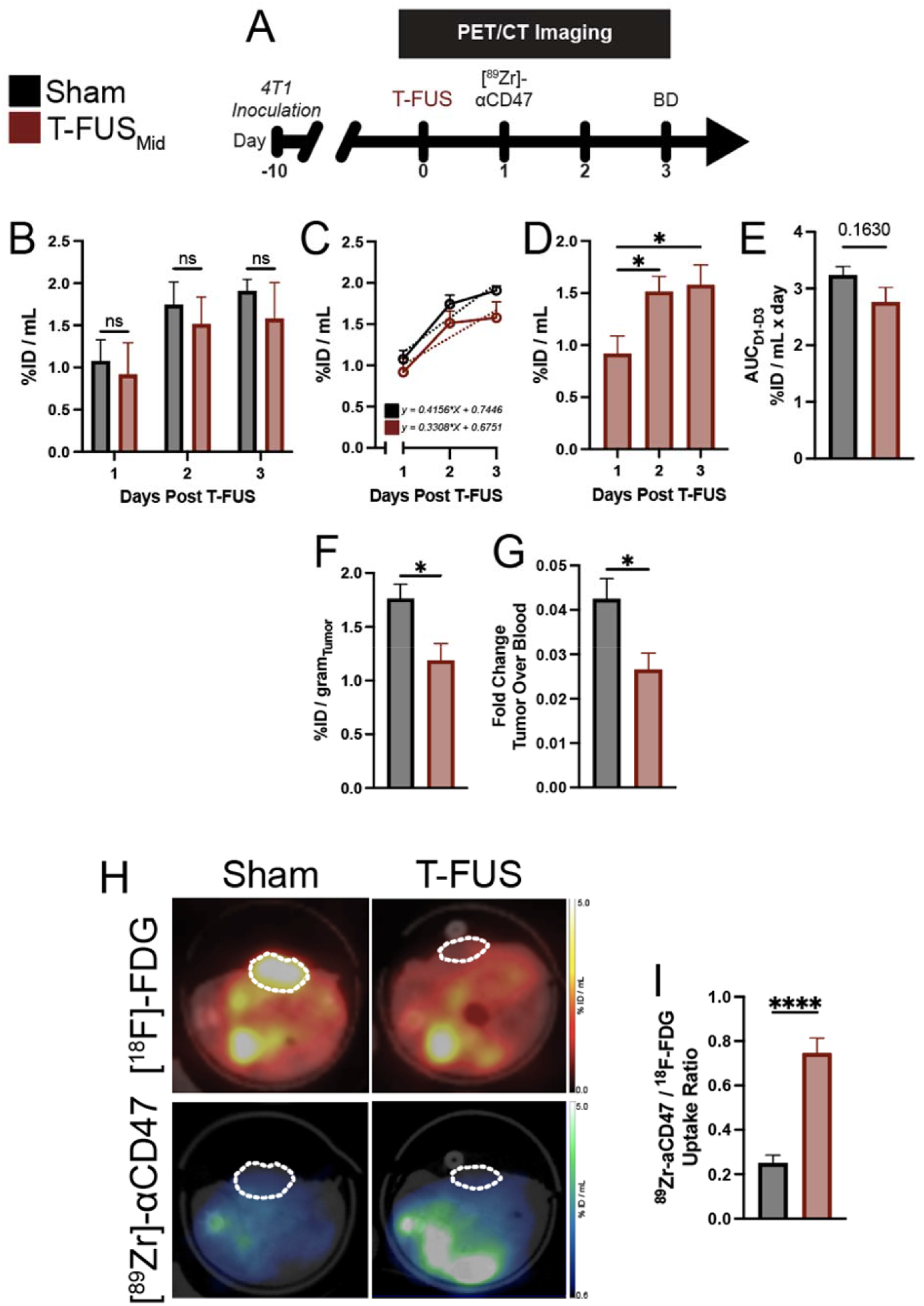
ImmunoPET reveals retained bulk antibody uptake following T-FUS_Mid_, with marked enrichment relative to residual viable tumor. A) Experimental timeline for T-FUS treatment, [^89^Zr]-αCD47 administration, serial PET/CT imaging, and endpoint biodistribution (BD) analysis. B) Longitudinal quantification of intratumoral [^89^Zr]-αCD47 uptake from PET/CT scans acquired on days 1-3 post-T-FUS. Significance assessed by two-way ANOVA. C) Linear regression analysis of longitudinal [^89^Zr]-αCD47 accumulation in sham and T-FUS-treated tumors. D) Quantification of intratumoral [^89^Zr]-αCD47 uptake in T-FUS-treated tumors over time, demonstrating increased antibody signal from day 1 to days 2 and 3 post-ablation. *p<0.05 vs indicated groups. E) Tumor-drug exposure in sham and T-FUS tumors, represented via area under the curve (AUC) analysis of intratumoral [^89^Zr]-αCD47 signal. Significance assessed by unpaired t-test. Sham vs T-FUS (p=0.1630). F) Endpoint *ex vivo* tumor accumulation of [^89^Zr]-αCD47 three days after T-FUS treatment. Significance assessed by unpaired t-test. Sham vs T-FUS_Mid_ (p = 0.0204). G) Tumor-o-blood ratio of [^89^Zr]-αCD47 uptake at endpoint. Significance assessed by unpaired t-test. Sham vs T-FUS_Mid_ (p = 0.0281). H) Representative tumor-matched [^18^F]-FDG and [^89^Zr]-αCD47 PET/CT images from sham and T-FUS mice. Dashed white outlines indicate tumor boundaries. I) Viable tumor-normalized antibody uptake, expressed as the ratio of [^89^Zr]-αCD47 to [^18^F]-FDG signal. Significance assessed by unpaired t-test. Sham vs T-FUS_Mid_ (p<0.0001). n = 5-6 per group.

Endpoint gamma counting revealed moderately reduced bulk [^89^Zr]-αCD47 uptake and tumor-to-blood [^89^Zr]-αCD47 signal ratio in T-FUS_Mid_-treated tumors compared to shams (Figure 8F,G). However, T-FUS_Mid_ did not alter circulating tracer levels or peripheral tissue distribution in the blood, heart, spleen, kidney, lung, or tumor-draining lymph node (Figure S3), indicating that the dominant effect was localized to the treated tumor rather than driven by systemic redistribution of the antibody. Because T-FUS_Mid_ also substantially reduced metabolically active tumor volume, these global uptake metrics likely reflect the smaller viable, perfused tumor compartment remaining after ablation rather than a simple failure of antibody delivery.

### Residual viable tumor is enriched for antibody uptake following Goldilocks T-FUS

ImmunoPET findings on bulk antibody uptake prompted a second analysis focused specifically on antibody accumulation within viable residual tumor. Because whole-tumor [^89^Zr]-αCD47 PET cannot distinguish ablated tissue from metabolically active tumor, we integrated the immunoPET readout with the matched [^18^F]-FDG signal to normalize antibody uptake to viable tumor fraction for each tumor (Figure 8H). Representative PET images showed reduced FDG-avid tumor volume after T-FUS_Mid_, with retained [^89^Zr]-αCD47 signal in residual tumor regions, suggesting that antibody uptake was concentrated within the remaining viable compartment.

Using this viable tumor-normalized metric, T-FUS_Mid_ significantly increased [^89^Zr]-αCD47 delivery relative to sham controls (Figure 8I). Specifically, after accounting for the reduction in FDG-defined viable tumor burden, T-FUS_Mid_ produced an approximately 3.2-fold increase in [^89^Zr]-αCD47 enrichment relative to residual tumor tissue – suggesting disproportionate penetrance and retention of antibody within the metabolically active residual compartment of ablated tumors. These findings indicate that Goldilocks T-FUS can achieve metabolic tumor debulking while preserving, and ultimately enriching, antibody exposure within viable peri-ablative tumor margins.

## DISCUSSION

The objective of this study was to define a therapeutically productive thermal dosing window - a “Goldilocks Zone” - for T-FUS that achieves meaningful tumor debulking while preserving the features needed to support local antibody access and activity. Using αCD47 as a model tumor-targeting immunotherapy, we developed for the first time a multimodal PET framework to evaluate the relationship between metabolic tumor debulking and immunotherapy delivery following thermal ablation. Although prior preclinical studies^9–11,21^ and emerging clinical data^6–8^ support the immunomodulatory potential of T-FUS, the thermal dose conditions that best balance cytoreduction, vascular preservation, and therapeutic access remain poorly defined. Here, *in silico* modeling established dose-escalated T-FUS regimens with graded increases in focal temperature and predicted ablation volume. *In vivo*, these regimens produced dose-dependent tumor debulking, broad hypoxia relief, vascular disruption, and thermal dose-sensitive impairment of functional perfusion. By integrating ultrasound-guided T-FUS, CEUS perfusion imaging, [^18^F]-FDG PET, and [^89^Zr]-αCD47 immunoPET, we identified an operational Goldilocks T-FUS condition that reduced metabolically active tumor burden while preserving antibody access and enriching αCD47 signal within viable residual tumor by approximately 3-fold. These findings establish thermal dose as a critical design parameter for T-FUS + immunotherapy combinations and provide a PET-informed framework for selecting ablative regimens that balance tumor destruction with therapeutic delivery. Taken together, this work offers critical, translationally relevant considerations for maximizing the therapeutic benefits of T-FUS treatment in both preclinical and clinical settings.

Thermal dose served as the organizing variable for this study because it links controllable FUS parameters to biological effect and is more readily mappable across systems. T-FUS exposures can be tuned through transducer geometry, frequency, acoustic power, sonication duration, and the number and spatial pattern of sonications^22–24^. However, the biological consequences of these parameters are ultimately governed by the magnitude and duration of tissue heating. For this reason, CEM43-based modeling provided a practical framework for generating three discrete thermal dose regimens - T-FUS_Low_, T-FUS_Mid_, and T-FUS_High_ - that could be compared at the levels of predicted temperature rise and ablation volume. These *in silico* studies confirmed that increasing input amplitude produced graded increases in maximum focal temperature, temperature-time exposure, and predicted ablated volume, thereby establishing a rational experimental basis for evaluating how thermal dose shapes the post-ablation tumor microenvironment. Importantly, TTC staining and H&E confirmed that the *in silico*-derived thermal doses yielded distinct, graded levels of tumor ablations *in vivo*, evidenced by tissue devitalization and coagulative necrosis within the targeted region. The ablative/periablative demarcations observed in this preclinical investigation directly mirror a number of FUS immuno-oncology clinical trial protocols (e.g. NCT06472661, NCT03237572, NCT04123535) in which a subtotal tumor volume of ∼30-50% is thermally ablated.

A provocative finding of this study was the reversal of tumor hypoxia across T-FUS conditions, without a strict relationship to thermal dose - suggesting that oxygenation-related changes after ablation reflect a more complex balance between tumor destruction, vascular injury, and peri-ablative tissue physiology. Indeed, broader literature shows that hyperthermic exposures can increase blood flow, improve oxygenation, and reduce hypoxia^25–30^. Whereas higher thermal doses can induce vascular occlusion, coagulative necrosis, and perfusion loss as demonstrated with other ablative modalities^31,32^, our findings suggest that T-FUS can shift the tumor microenvironment toward reduced hypoxia when applied within an appropriate thermal range. These observations are also consistent with previously published work demonstrating that FUS ablation, albeit mechanical (histotripsy), is capable of abrogating tumor hypoxia^33^.

In this study, endomucin staining and CEUS perfusion imaging provided complementary, though not fully independent, readouts of the post-ablation vascular state. Reduced endomucin-positive vascular area and diminished contrast wash-in may reflect direct vascular injury as well as acute post-thermal tissue effects, including edema or compression of perfused microvessels. Of note, due to measurement limitations, bulk signal reduction may have diluted any relative enhancements in blood flow within peri-ablative regions. Our findings support the central premise that thermal ablation generates competing biological effects. While sufficient heating is needed for cytoreduction, excessive vascular disruption may limit access to residual viable tumor by macromolecular immune therapeutics or immune cells^34–36^. This tradeoff generated the rationale for selecting T-FUS_Mid_ for subsequent PET-based studies. T-FUS_Low_ produced measurable biological effects but did not maximize tumor debulking. By contrast, T-FUS_High_ produced more extensive thermal injury but was accompanied by a disproportionate reduction in functional perfusion, raising concern that near-complete ablation could compromise the physiological features required for post-ablation therapeutic delivery. T-FUS_Mid_ therefore represented a foray into the postulated “Goldilocks Zone” by achieving robust partial ablation and marked hypoxia relief while avoiding the severe perfusion deficit observed at the highest thermal dose. In this context, the “Goldilocks Zone” should not be interpreted as an optimum, but rather as a candidate physiologic thermal window in which tumor cytoreduction is achieved without fully eliminating the residual vascular and stromal features needed to support therapeutic access.

Quantitative PET imaging was deployed as a non-terminal modality for volumetric assessment of metabolic activity following T-FUS_Mid_ treatment. Indeed, metabolic PET imaging provides a critical advantage over traditional gross staining and histology for quantitative, volumetric, spatiotemporally resolved assessment of ablation. Consistent with our histological findings and other clinical studies deploying [^18^F]-FDG PET/CT imaging following ablative procedures^18,37^, we observed a marked reduction in FDG signal within ablated tumors – suggesting that TTC staining and H&E may underestimate the full extent of metabolically compromised tumor tissue after T-FUS_Mid_. To our knowledge, this study represents the first preclinical application of dynamic FDG PET imaging to assess ablation following T-FUS treatment, yielding opportunities and insights relevant to modern clinical practice.

A primary goal of this study was to determine whether partial T-FUS remains permissive to the delivery of large biologics, including tumor-targeted antibodies. Prior work has shown that FUS thermal ablation can enhance local accumulation of systemically administered agents in mammary carcinoma models, including small-molecule MRI contrast agents and long-circulating liposomal drug carriers^38,39^. To this end, ablation has been shown to increase liposomal and free chemotherapy accumulation within treated tumors, with prominent accumulation in the viable rim surrounding ablated tissue. Here, we extend this principle to antibody-scale biologics for the first time using immunoPET imaging, wherein αCD47 was selected as a representative tumor-targeted antibody. CD47 is enriched in 4T1 tumors^40^ and is relevant to the biology of breast cancer^41,42^ and other solid tumor types to which these findings could extend. The objective of this study was not to test CD47 blockade as a therapeutic intervention, but to use radiolabeled αCD47 as a model payload for quantifying antibody access after thermal ablation. Strikingly, serial PET/CT imaging revealed that intratumoral [^89^Zr]-αCD47 signal was retained within T-FUS-treated tumors and increased over the longitudinal imaging window at a rate comparable to shams. These data indicate that partial thermal ablation, as practiced in FUS immuno-oncology paradigms, is not uniformly deleterious to localized immunotherapeutic access and may instead preserve a targetable residual tumor niche for large biologics when delivered within an appropriate thermal window.

The apparent discrepancy between retained *in vivo* PET signal and reduced endpoint bulk uptake underscores the importance of interpreting antibody delivery in relation to residual viable tumor burden after ablation. Whole-tumor antibody uptake metrics cannot distinguish nonviable ablated tissue from metabolically active tumor and may therefore underestimate delivery efficiency when effective cytoreduction has occurred. To address this, we integrated the [^89^Zr]-αCD47 immunoPET with tumor-matched [^18^F]-FDG PET to normalize antibody uptake to metabolically active residual tumor. Using this approach, T-FUS_Mid_ produced a greater than 3-fold increase in αCD47 enrichment relative to sham controls. Thus, T-FUS can not only preserve antibody access, but may concentrate antibody exposure within the remaining viable tumor compartment. While PET resolution was insufficient to definitively localize antibody signal to periablative margins, these findings are consistent with reports that hyperthermia can improve antibody delivery in solid tumor settings^29,43–48^.

Several focused questions remain important for future investigation. Spatially resolved analyses will be important to determine whether antibody accumulation occurs preferentially within periablative margins and whether this enrichment reflects changes in perfusion, vascular permeability, target availability, stromal remodeling, immune remodeling, or some combination of these mechanisms. These studies will be particularly important for defining how thermal dose influences not only antibody delivery, but also the immunologic consequences of T-FUS. Future studies should also determine whether this delivery advantage extends across additional tumor models, antibody platforms, and treatment schedules. Indeed, some therapeutic studies implement neoadjuvant treatment schema^10,11,14^ while others deploy adjuvant dosing^49–51^. Despite the lack of consensus on optimal timing of therapeutic administration relative to ablation, timing remains a critical determinant of synergy, and these findings underscore the need to further dissect the interplay between thermal dose and therapeutic timing.

Taken together, these findings support thermal dose as a key variable for cooperative T-FUS immunotherapy combinations. We highlight the importance and availability of treatment parameters that debulk tumor while preserving an immunotherapy-accessible residual tumor niche, herein focusing on thermal dose. We highlight the power of a multimodal imaging framework for non-invasively and longitudinally evaluating the tradeoffs between localized thermal destruction and therapeutic access. As ablative modalities like T-FUS continue to advance in immuno-oncology, PET-informed strategies such as this may help rationally define and curate ablative regimens that drive local tumor control while enabling effective delivery of immune-directed therapies.

## MATERIALS & METHODS

### In silico Focused Ultrasound Modeling

We model linear acoustic propagation with absorption and heterogeneous background properties using the coupled first-order system solved by k-Wave^52,53^:

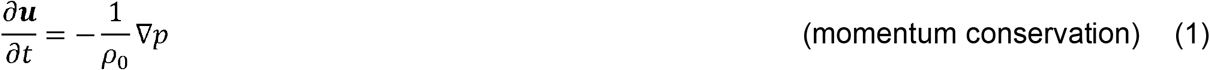

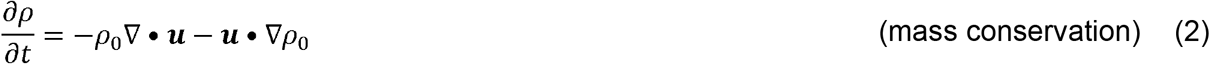

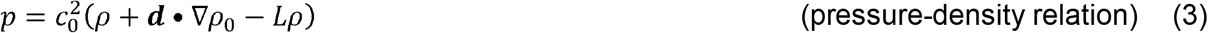

Where *u* is the acoustic particle velocity, *p* is the acoustic pressure, *ρ* is the acoustic density, *ρ*_0_ is ambient (or equilibrium) density, *c*_0_ is the isentropic sound speed, *d* is the acoustic particle displacement, and the operator *L* in the pressure-density relation is a linear integro-differential operator that accounts for acoustic absorption and dispersion that follows a frequency power law.

The temperature field was obtained by solving the Pennes’ bioheat equation^54^:

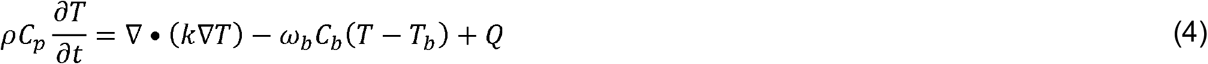

Where T is temperature, *C*_*p*_ is the specific heat capacity, *k* is the thermal conductivity, is *ω*_*b*_ blood perfusion rate, *C*_*b*_ and *T*_*b*_ are blood specific heat and arterial temperature, and *Q* is the volumetric heat source.

In this model, the heat source *Q* corresponds to the acoustic power absorbed per unit volume due to ultrasound propagation. This was computed from the simulated acoustic field as^55^:

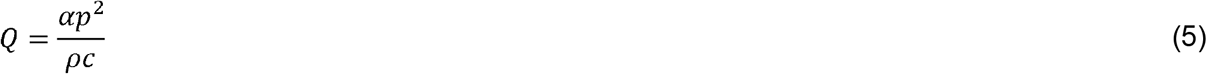

Where *α* is the acoustic absorption coefficient.

The concept of thermal dose is based on the relationship between tissue heating, exposure time, and biological response. The CEM43 is defined as^56^:

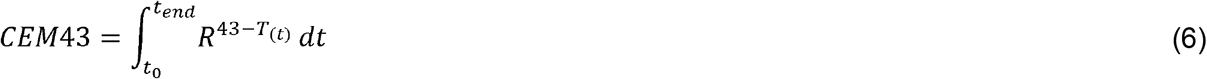

Where *t*_0_ and *t*_*end*_ are the start and end times of the sonication,*T*_*(t)*_ is the transient tissue temperature, and is the iso-effect constant, with *R* =0.5 for > 43 °C and 0.25 for *T* < 43 °C. Once the accumulated dose exceeds a defined threshold, the tissue is considered irreversibly damaged. A commonly accepted threshold for coagulative necrosis is *CEM*43 > 240 minutes^57^, which is adopted here as the criterion for ablation.

A 3D computational model was developed to simulate focused ultrasound (FUS) ablation of tumor tissue. The geometry consisted of layered domains representing skin and underlying tumor tissue, with dimensions chosen to approximate the experimental configuration. A schematic of the computational domain is shown in Figure S1A.

For acoustic propagation, the k-Wave MATLAB toolbox was employed. The model solved the coupled first-order acoustic equations under the assumption of linear wave propagation. A perfectly matched layer (PML) was applied to the boundaries of the computational grid to eliminate artificial reflections. Hydrophone measurements were used to calibrate the input pressure amplitude in the model. The driving parameters and the corresponding simulated focal peak-to-peak (P-P) pressures within tumor tissue are summarized in Table S3.

The resulting acoustic intensity distribution was used to compute the absorbed acoustic power per unit volume, which served as the heat source in the Pennes bioheat equation. Tumor ablation was quantified using the CEM43 thermal dose metric. All simulations were run on a 620 × 461 × 314 grid with a spatial step size of 170 μm, giving a maximum supported frequency of 4.36 MHz. The temporal step size was 33 ns, corresponding to a Courant–Friedrichs–Lewy (CFL) number of 0.3, and acoustic simulations were run for 2841-time steps. For the thermal simulations, a time step size of 0.1 s was used. The initial tissue temperature was set to 37 °C, and the blood temperature *T*_*b*_ was also assumed to be 37 °C. To confirm numerical stability, a grid independence study was performed. The peak-to-peak pressure at the acoustic focus was compared across grids of varying resolution, and the results are shown in Figure S1B. Table S2 summarizes the material properties assigned to each tissue layer in this study.

### Cell line and culture

Triple-negative 4T1 murine breast cancer cells were purchased from ATCC (ATCC CRL-2539). 4T1 cells were maintained at 37ºC and 5% *Co*_2_ and cultured in RPMI 1640 (+ L glut, Gibco #11875-093) supplemented with 10% Fetal Bovine Serum (FBS). Upon thawing procedure, cells were passaged up to a maximum of three times and expanded in a logarithmic growth phase prior to implantation. In advance of expansion and storage, 4T1 cells were authenticated and screened for mycoplasma.

### Syngeneic mouse tumor model

All murine experiments were approved and performed in accordance with the guidelines and regulations of the University of Virginia and the University of Virginia Animal Care and Use Committee. Female BALB/c mice were purchased from The Jackson Laboratory (Jax # 000651) between 6-8 weeks of age. Mice were housed in pathogen-free animal facility within a stable environment and had access to food and water at all times. Mice were anesthetized with intraperitoneal (i.p.) injection of ketamine (50 mg/kg; Zoetis) and dexdomitor (0.25 mg/kg; Dechra) in 0.9% saline (ket/dex). Following, mice were shaved on the right flank and subcutaneously (s.c.) implanted with 4*10^5^ 4T1 cells. Mice were weighed weekly, and tumors were monitored daily following T-FUS treatment with precision digital caliper. Digital caliper measurements were used to calculate tumor volume as follows: *volume* (*length × widht^2^*) /2. Mice were randomized into treatment groups using an in-house algorithm designed to minimize standard deviation in starting average tumor volume among groups.

### Ultrasound-guided FUS partial thermal ablation

On the day of T-FUS treatment, mice were anesthetized with an i.p. ket/dex, and tumors were depilated prior to the procedure. Thermal ablation was performed with a custom-built ultrasound-guided FUS system comprised of four 3.78 MHz single-element transducers (SU-102, Sonic Concepts), each with a diameter of 33 mm and a radius of curvature of 55 mm. The system was powered by a 200W amplifier (Electronics & Innovation 1020L) driven by an arbitrary function generator (Tektronix AFG3022C), and axially co-registered to a 15 MHz MicroScan linear ultrasound imaging array (MS200; Visual Sonics). Degassed deionized water, maintained at 37ºC, was used as the acoustic coupling medium. Tumor positioning was achieved via a motorized 3D motion stage. Tumor localization was achieved using B-mode imaging (VEVO 2100 MS200). T-FUS_Low_, was applied continuously for 20 s at 9.8 W acoustic power, while T-FUS_Mid_ and T-FUS_High_ were applied for 15s at 18 W and 28.7 W acoustic power, respectively. Thermal ablation was carried out over 2-3 ultrasound-visible tumor planes, each separated by 1 mm. Each plane underwent 2-5 sonications separated by 1.5mm spacing and scaled based on tumor volume. Mice that did not receive T-FUS treatment underwent “sham” treatment, consisting of anesthesia, depilation, and exposure to the 37ºC degassed water bath for 3 minutes. At the conclusion of “sham” or T-FUS treatments, mice were transferred to a heating pad, given Antisedan for anesthesia reversal, and recovered.

### Contrast-enhanced Ultrasound Perfusion Imaging

Contrast-enhanced ultrasound (CEUS) perfusion imaging of T-FUS-treated and Sham control tumors was performed immediately following thermal ablation for 20 seconds. Lipid-shelled decafluorbutane microbubbles (MB) were infused at 1 × 10^7^ min^−1^ (25uL/min), prepared in accordance with previously published methods^58^, and imaged using the eL18-4 linear array and Philips EPIQ Elite System. During infusion, ultrasound images were acquired at a frame rate of 93Hz following a brief high-mechanical index sequence of 0.93 to effectively destroy MBs. The Philips QLAB Quantification Software was used to generate time-intensity curves from CEUS data, enabling quantification of MBs wash-in slope or flux rate (dB/s) and peak intensity (dB). Two independent video acquisitions were analyzed per tumor, and the resulting measurements were averaged to obtain a single representative value for each tumor.

### Pimonidazole Injections

Pimonidazole (Hypoxyprobe) was diluted at a concentration of 60mg/kg in sterile PBS and administered through via i.p. injection. Mice were sacrificed 1.5 hours after injection.

### Immunohistochemistry

On day 13 post-inoculation, Sham and T-FUS treated 4T1-treated mice underwent cardiac perfusion with a sequence of 1X PBS, 10% neutral buffered formalin (NBF), 1X PBS followed by tumor excision. 24 hours later, fixed tumors were paraffin embedded, sectioned, and stained for hematoxylin and eosin (H&E) or 2,3,5-Triphenyltetrazolium chloride (TTC). Digital scans of H&E stained slides were collected using ZEISS Axioscan 7.

### Immunofluorescence Imaging

For immunofluorescence staining of FFPE sections, slides were deparaffinized in a series of xylene and alcohol solutions. To retrieve antigens, slides were placed in sodium citrate solution and heated in a pressure cooker on the high setting for 25 minutes. Slides were then permeabilized with 0.1% Triton-X in PBS for 15 minutes at room temperature, washed in 1x PBS, then placed in 0.25% (w/v) Sudan Black B in 70% isopropyl alcohol for 1.5 hours. Tissue sections were then blocked with donkey serum for 1 hour at room temperature and primary antibodies were incubated with tissue sections overnight at 4C. Primary antibodies were diluted in antibody buffer (5% donkey serum and 1% BSA in 1x PBS). The primary antibodies used were anti-pimonidazole (1:200, PAB2627) and anti-endomucin (1:100, sc-65495). Slides were mounted in 50/50 glycerol and coverslipped. Sections were imaged on the Leica Thunder with a 20x objective. For vessel analysis, AngioTool^59^ was used to quantify vascular characteristics. For hypoxia analysis, anti-pimonidazole-positive area coverage was quantified following image thresholding in Fiji (ImageJ).

### Whole-body PET/CT Imaging and Analysis

Dynamic or static PET/CT imaging scans were performed at select time points between 0- and 3-days following T-FUS treatment. Mice were anesthetized with isoflurane/oxygen gas mixture (2% for induction) using a vaporizer. Tumor-bearing mice were i.v. injected with 150 μCi per mouse^60^ of ^18^F-Fludeoxyglucose (FDG) 4-hours after thermal ablation. Concomitant with administration of ^18^F-FDG, mice underwent 60-minute list mode (LM) whole-body dynamic positron emission tomography (PET) acquisition using a similar protocol as described^61,62^, followed by a 10-minute computed tomography (CT) scan (250 projections, 5 min approximate exposure time, 500 μM voxel size) using an Albira Si small-animal trimodal PET/SPECT/CT system (Bruker, Billerica, MA). Specifically for the dynamic PET scan, the LM data were then sorted into 23 time bins, with frames x times (s) as follows: (11 × 8s; 1 × 12s; 2 × 60s; 1 × 180s; 8 × 400s). One day following T-FUS treatment, mice received 100 μCi of ^89^Zr-αCD47 and underwent a 20-minute static PET and 20-minute CT scan (600 projections, 10-12 min approximate exposure time, 125 μM voxel size). Repeat PET and CT imaging scans were performed on days 2 and 3 following thermal ablation. Reconstruction was performed on dynamic and static PET images uses the Maximum Likelihood Expectation Maximization (MLEM) 0.5mm algorithm, with a total of 12 iteration and incorporation scatter, decay, and random corrections. CT scans were reconstructed using a 3D filtered back projection (FBP) algorithm and high resolution. Co-registration and analysis of PET and CT scans was performed using PMOD 3.9 (PMOD Technologies Ltd.). Standard tumor uptake values were determined by volume-of-interest (VOI) analysis, with tumor boundaries delineated using anatomical landmarks from corresponding high-resolution CT images.

### Ex Vivo Biodistribution Analysis

On day 3 following treatment, mice receiving i.v. administration of [^89^Zr]-αCD47 were euthanized, and peripheral organs and tumors were collected for biodistribution analysis. Tumor, heart, spleen, blood, tumor-draining lymph node (inguinal), kidney, and lungs were excised. Tissue samples were transferred to a pre-weighed scintillation vial and analyzed using an automatic gamma counter (Hidex). The percent injected dose per gram (%ID/g) was calculated after correcting for background and radioactive decay and was compared to an aqueous [□ □Zr] standard.

### ^18^F-FDG Production and Purification

^18^F-FDG is commercially available from PETNET Solutions Inc (Siemens Healthineers, Charlottesville, VA) and was used without subsequent purification for all studies.

### Conjugation and Radiolabeling of ^89^Zr-αCD47

Using a 30K molecular weight cut-off centrifugal filter, 5mg αCD47 (BioXCell, MIAP410) was exchanged to 0.1M Na2CO3-NaHCO3 buffer pH 9.0 and final volume was adjusted to 1mL by adding the same buffer. Separately, 1.3mg of p-isothiocynate-benzyl-DFO was dissolved in 208uL of DMSO to prepare a stock solution. A 20 uL aliquot of the DFO solution was added to 1mL solution containing 5mg of αCD47. The mixture was incubated for 45 minutes at 37° C. Following incubation, the mixture underwent purification with a PD10 gel filtration column, eluting with 0.25M sodium acetate solution pH 6.0. Incubation of the mixture with 5uL (3 mCi) of ^89^Zr oxalate occurred for 1 hour. Subsequently, the mixture underwent gel filtration purification with a PD10 column, eluting with 0.9% saline for labeling yield and purity confirmed by size-exclusion HPLC (Superdex 200 Increase 5/150 GL column with 0.45ml/min flow rate of water elution).

### Statistical analysis

Statistical analyses were performed using GraphPad Prism 10.1.0. A comprehensive overview of the statistical techniques used in each experiment can be found in the accompanying figure legend. “n.s.” denotes that data are not significant. All data are represented as mean ± SEM.

## Supporting information

Demir-Goldilocks-Supplement

## Acknowledgments

We thank Dr. Frederic Padilla for his expertise and development of the ultrasound-guided FUS system used herein. This research was supported by state funding within the University of Virginia Comprehensive Cancer Center and NCI Cancer Center Support Grant P30CA44579. This work was supported by Biorepository and Tissue Research Facility which is supported by the University of Virginia School of Medicine, Research Resource Identifiers (RRID): SCR_022971. This work used FFPE embedded and sectioning in the Research Histology Core Facility which is supported by the University of Virginia School of Medicine, Research Resource Identifiers (RRID): SCR_025470. We thank Megan Burts and Dr. Maurits Jansen for their expertise and support with PET/CT scans. This work used the Bruker Albira Si small-animal trimodal PET/SPECT/CT system in the Molecular Imaging Core Facility which is supported by the University of Virginia School of Medicine, Research Resource Identifiers (RRID): SCR_025472. Imaging data was acquired through the University of Virginia Molecular Imaging Core Lab, with NIH S10OD025024 funding for the Bruker 9.4T MRI scanner. We thank Dr. Damodara Naidu Kommi and Dr. Shivashankar Khanapur for their radiochemistry assistance and support. This work used the Hidex Automatic Gamma Counter and [^18^F]-FDG supplied by the Radiochemistry Core Facility, which is supported by the University of Virginia School of Medicine, Research Resource Identifiers (RRID): SCR_025472 and SCR_025471.

## Funding

Supported by NIH Director’s Early Independent Award (DP5OD031846), US DOD BCRP Era of Hope Scholar Award (HT9425-25-1-0411), Wallace H. Coulter Foundation for Translational Research Grant to NDS; UVA Focused Ultrasound Immuno-Oncology (FUSION) Center pilot award to SK and NDS; and NCI U01CA243007 to MJL. ZEFD and TS were supported by UVA Cancer Biology Training Grant (NIH T32CA009109). ZEFD was additionally supported by the UVA Cancer Center Training Fellowship and UVA SEAS Endowed Victor Orphan Graduate Fellowship.

## Author Contributions

Conceptualization: ZEFD, TS, NDS

Methodology: ZEFD, TS, MRD, PT, CPG, MAJ, DNK, SK, ALK, SMP, MJL, JRL, JH, BK, NDS

Formal Analysis: ZEFD, TS, PT, MRD, CPG

Investigation: ZEFD, TS, MRD, PT, MRB, MAJ, DNK, SK,

Writing - Original Draft: ZEFD

Writing - Review and Editing: ZEFD, TS, MRD, KDN, SMP, MJL, BK, NDS

Resources: MRB, MAJ, DNK, SK, CPG, SMP, MJL

Funding acquisition: NDS

Supervision: NDS

## Notes

### Competing Interest Statement

The authors have declared no competing interest.

## REFERENCES

1. Wang, L.-H., Jiang, Y., Sun, C.-H.Chen, P.-T. & Ding, Y.-N. Advancements in the application of ablative therapy and its combination with immunotherapy in anti-cancer therapy. Biochimica et Biophysica Acta (BBA) -Reviews on Cancer 1880, 189285 (2025).

2. Carriero, S. et al. Ablative Therapies for Breast Cancer: State of Art. Technol Cancer Res Treat 22, 15330338231157193 (2023).

3. Granata, V. et al. Local ablation of pancreatic tumors: State of the art and future perspectives. World J Gastroenterol 27, 3413–3428 (2021).

4. Demir, Z. E. F. & Sheybani, N. D. Therapeutic Ultrasound for Multimodal Cancer Treatment: A Spotlight on Breast Cancer. Annual Review of Biomedical Engineering 27, 371–402 (2025).

5. DeWitt, M., Demir, Z. E. F., Sherlock, T., Brenin, D. R. & Sheybani, N. D. MR Imaging-Guided Focused Ultrasound for Breast Tumors. Magnetic Resonance Imaging Clinics 0, (2024).

6. Zhu, X.-Q. et al. Alterations in Immune Response Profile of Tumor-Draining Lymph Nodes after High-Intensity Focused Ultrasound Ablation of Breast Cancer Patients. Cells 10, 3346 (2021).

7. Lu, P. et al. Increased infiltration of activated tumor-infiltrating lymphocytes after high intensity focused ultrasound ablation of human breast cancer. Surgery 145, 286–293 (2009).

8. Xu, Z.-L. et al. Activation of tumor-infiltrating antigen presenting cells by high intensity focused ultrasound ablation of human breast cancer. Ultrasound Med Biol 35, 50–57 (2009).

9. Sheybani, N. D. et al. Combination of thermally ablative focused ultrasound with gemcitabine controls breast cancer via adaptive immunity. J Immunother Cancer 8, e001008 (2020).

10. Silvestrini, M. T. et al. Priming is key to effective incorporation of image-guided thermal ablation into immunotherapy protocols. JCI Insight 2, e90521 (2017).

11. Fite, B. Z. et al. Immune modulation resulting from MR-guided high intensity focused ultrasound in a model of murine breast cancer. Sci Rep 11, 927 (2021).

12. Wang, J. et al. Multiomic analysis for optimization of combined focal and immunotherapy protocols in murine pancreatic cancer. Theranostics 12, 7884–7902 (2022).

13. Do, H. D. et al. Combination of thermal ablation by focused ultrasound, pFAR4-IL-12 transfection and lipidic adjuvant provide a distal immune response. Explor Target Antitumor Ther 3, 398–413 (2022).

14. Chavez, M. et al. Distinct immune signatures in directly treated and distant tumors result from TLR adjuvants and focal ablation. Theranostics 8, 3611–3628 (2018).

15. Liu, F. et al. Boosting high-intensity focused ultrasound-induced anti-tumor immunity using a sparse-scan strategy that can more effectively promote dendritic cell maturation. J Transl Med 8, 7 (2010).

16. Hu, Z. et al. Investigation of HIFU-induced anti-tumor immunity in a murine tumor model. J Transl Med 5, 34 (2007).

17. Xing, Y., Lu, X., Pua, E. C. & Zhong, P. The effect of high intensity focused ultrasound treatment on metastases in a murine melanoma model. Biochemical and Biophysical Research Communications 375, 645–650 (2008).

18. Aarntzen, E. H. J. G., Heijmen, L. & Oyen, W. J. G. 18F-FDG PET/CT in Local Ablative Therapies: A Systematic Review. Journal of Nuclear Medicine 59, 551–556 (2018).

19. Orsi, F. et al. High-Intensity Focused Ultrasound Ablation: Effective and Safe Therapy for Solid Tumors in Difficult Locations. American Journal of Roentgenology 195, W245–W252 (2010).

20. Bongiovanni, A. et al. 3-T magnetic resonance–guided high-intensity focused ultrasound (3 T-MR-HIFU) for the treatment of pain from bone metastases of solid tumors. Support Care Cancer 30, 5737–5745 (2022).

21. Deng, J., Zhang, Y., Feng, J. & Wu, F. Dendritic Cells Loaded with Ultrasound-Ablated Tumour Induce in vivo Specific Antitumour Immune Responses. Ultrasound in Medicine and Biology 36, 441–448 (2010).

22. Moussa, M. et al. Effect of thermal dose on heat shock protein expression after radio-frequency ablation with and without adjuvant nanoparticle chemotherapies. International Journal of Hyperthermia 32, 829–841 (2016).

23. Bie, K. C. C. de et al. Outcomes of CEM43 in Predicting Thermal Damage Induced by Focal Laser Ablation in Controlled Ex Vivo Experiments: A Comparison to Histology and MRI. Lasers in Surgery and Medicine 56, 723–733 (2024).

24. Jones, R. M. et al. Accumulated thermal dose in MRI-guided focused ultrasound for essential tremor: repeated sonications with low focal temperatures. J Neurosurg 132, 1802–1809 (2020).

25. Hannon, G., Tansi, F. L., Hilger, I. & Prina-Mello, A. The Effects of Localized Heat on the Hallmarks of Cancer. Advanced Therapeutics 4, 2000267 (2021).

26. Bosque, J. J., Calvo, G. F., Pérez-García, V. M. & Navarro, M. C. The interplay of blood flow and temperature in regional hyperthermia: a mathematical approach. R Soc Open Sci 8, 201234 (2021).

27. Kong, G., Braun, R. D. & Dewhirst, M. W. Hyperthermia Enables Tumor-specific Nanoparticle Delivery: Effect of Particle Size1. Cancer Res 60, 4440–4445 (2000).

28. Karino, T., Koga, S. & Maeta, M. Experimental studies of the effects of local hyperthermia on blood flow, oxygen pressure and pH in tumors. The Japanese Journal of Surgery 18, 276–283 (1988).

29. Sun, X., Xing, L., Clifton Ling, C. & Li, G. C. The effect of mild temperature hyperthermia on tumour hypoxia and blood perfusion: relevance for radiotherapy, vascular targeting and imaging. International Journal of Hyperthermia 26, 224–231 (2010).

30. Centelles, M. N., Wright, M., Gedroyc, W. & Thanou, M. Focused ultrasound induced hyperthermia accelerates and increases the uptake of anti-HER-2 antibodies in a xenograft model. Pharmacological Research 114, 144–151 (2016).

31. Moussa, M. et al. Radiofrequency Ablation–Induced Upregulation of Hypoxia-Inducible Factor-1α Can Be Suppressed with Adjuvant Bortezomib or Liposomal Chemotherapy. Journal of Vascular and Interventional Radiology 25, 1972–1982 (2014).

32. Tong, Y. et al. Effect of a hypoxic microenvironment after radiofrequency ablation on residual hepatocellular cell migration and invasion. Cancer Science 108, 753–762 (2017).

33. Song, B. et al. Histotripsy-focused ultrasound treatment abrogates tumor hypoxia responses and stimulates anti-tumor immune responses in melanoma. Mol Cancer Ther 24, 1088–1098 (2025).

34. Melssen, M. M., Sheybani, N. D., Leick, K. M. & Slingluff, C. L. Barriers to immune cell infiltration in tumors. J Immunother Cancer 11, (2023).

35. Bordeau, B. M. & Balthasar, J. P. Strategies to enhance monoclonal antibody uptake and distribution in solid tumors. Cancer Biol Med 18, 649–664 (2021).

36. Tong, R. T. et al. Vascular Normalization by Vascular Endothelial Growth Factor Receptor 2 Blockade Induces a Pressure Gradient Across the Vasculature and Improves Drug Penetration in Tumors. Cancer Res 64, 3731–3736 (2004).

37. Orgera, G. et al. High-intensity focused ultrasound (HIFU) in patients with solid malignancies: evaluation of feasibility, local tumour response and clinical results. Radiol med 116, 734–748 (2011).

38. Wong, A. W. et al. Ultrasound ablation enhances drug accumulation and survival in mammary carcinoma models. J Clin Invest 126, 99–111 (2016).

39. Kim, H. R. et al. MRI Monitoring of Tumor-Selective Anticancer Drug Delivery with Stable Thermosensitive Liposomes Triggered by High-Intensity Focused Ultrasound. Mol. Pharmaceutics 13, 1528–1539 (2016).

40. Lian, S. et al. Dual blockage of both PD-L1 and CD47 enhances immunotherapy against circulating tumor cells. Sci Rep 9, 4532 (2019).

41. Yuan, J. et al. Combined high expression of CD47 and CD68 is a novel prognostic factor for breast cancer patients. Cancer Cell Int 19, 238 (2019).

42. Betancur, P. A. et al. A CD47-associated super-enhancer links pro-inflammatory signalling to CD47 upregulation in breast cancer. Nat Commun 8, 14802 (2017).

43. Hauck, M. L. et al. A local hyperthermia treatment which enhances antibody uptake in a glioma xenograft model does not affect tumour interstitial fluid pressure. Int J Hyperthermia 13, 307–316 (1997).

44. Schuster, J. M. et al. Hyperthermic modulation of radiolabelled antibody uptake in a human glioma xenograft and normal tissues. Int J Hyperthermia 11, 59–72 (1995).

45. Hosono, M. N., Hosono, M., Endo, K., Ueda, R. & Onoyama, Y. Effect of hyperthermia on tumor uptake of radiolabeled anti-neural cell adhesion molecule antibody in small-cell lung cancer xenografts. J Nucl Med 35, 504–509 (1994).

46. Hauck, M. L. & Zalutsky, M. R. Enhanced tumour uptake of radiolabelled antibodies by hyperthermia: Part I: Timing of injection relative to hyperthermia. International Journal of Hyperthermia 21, 1–11 (2005).

47. Farr, N. et al. Hyperthermia-enhanced targeted drug delivery using magnetic resonance-guided focussed ultrasound: a pre-clinical study in a genetic model of pancreatic cancer. Int J Hyperthermia 34, 284–291 (2018).

48. Frazier, N. & Ghandehari, H. Hyperthermia approaches for enhanced delivery of nanomedicines to solid tumors. Biotechnology and Bioengineering 112, 1967–1983 (2015).

49. Shi, L. et al. PD-1 Blockade Boosts Radiofrequency Ablation–Elicited Adaptive Immune Responses against Tumor.Clin Cancer Res 22, 1173–1184 (2016).

50. Jiang, M., Shao, Q., Slaughter, J. & Bischof, J. Irreversible Electroporation has More Synergistic Effect with Anti-PD-1 Immunotherapy than Thermal Ablation or Cryoablation, in a Colorectal Cancer Model. Advanced Therapeutics 7, 2400068 (2024).

51. Guo, R.-Q., Peng, J.-Z.Li, Y.-M. & Li, X.-G. Microwave ablation combined with anti-PD-1/CTLA-4 therapy induces an antitumor immune response to renal cell carcinoma in a murine model. Cell Cycle 22, 242–254 (2023).

52. Brown, D. G. & Insana, M. F. Acoustic scattering theories applied to biological tissues. The Journal of the Acoustical Society of America 91, 2406–2406 (1992).

53. Pierce, A. D. Mathematical Theory of Wave Propagation. in Encyclopedia of Acoustics (ed. Crocker, M. J.) 21–37 (Wiley, 1997). doi:10.1002/9780470172513.ch2.

54. Pennes, H. H. Analysis of Tissue and Arterid Blood Temperatures in the Resting Human Forearm. Journal of Applied Physiology 1, 93–122 (1948).

55. Wootton, J. H., Ross, A. B. & Diederich, C. J. Prostate thermal therapy with high intensity transurethral ultrasound: The impact of pelvic bone heating on treatment delivery. International Journal of Hyperthermia 23, 609–622 (2007).

56. Sapareto, S. A. & Dewey, W. C. Thermal dose determination in cancer therapy. International Journal of Radiation Oncology*Biology*Physics 10, 787–800 (1984).

57. Harris, G. R., Herman, B. A. & Myers, M. R. A Comparison of the Thermal-Dose Equation and the Intensity-Time Product, Itm, for Predicting Tissue Damage Thresholds. Ultrasound in Medicine & Biology 37, 580–586 (2011).

58. Belcik, J. T. et al. Augmentation of Muscle Blood Flow by Ultrasound Cavitation Is Mediated by ATP and Purinergic Signaling. Circulation 135, 1240–1252 (2017).

59. Zudaire, E., Gambardella, L., Kurcz, C. & Vermeren, S. A Computational Tool for Quantitative Analysis of Vascular Networks. PLoS ONE 6, e27385 (2011).

60. Molinos, C. et al. Low-Dose Imaging in a New Preclinical Total-Body PET/CT Scanner. Front. Med. 6, 88 (2019).

61. Zhong, M. & Kundu, B. K. Optimization of a Model Corrected Blood Input Function From Dynamic FDG-PET Images of Small Animal Heart In Vivo. IEEE Trans. Nucl. Sci. 60, 3417–3422 (2013).

62. Massey, J. C. et al. Model Corrected Blood Input Function to Compute Cerebral FDG Uptake Rates From Dynamic Total-Body PET Images of Rats in vivo. Front. Med. 8, 618645 (2021).

