## Supplementary material for "Multimodal PET Defines a ‘Goldilocks’ Thermal Window for Focused Ultrasound Ablation and Immunotherapy Combinations": Demir-Goldilocks-Supplement

<u>**Supplemental Materials For:**</u>

**Supplemental Table 1. Average tumor dimension and skin thickness.**

| Tumor Dimension (mm) | Skin Thickness (mm) |
|---|---|
| L = 5.94<br>W = 3.49 | 0.17 |

Supplemental Table 2. Acoustic and thermophysical properties of the materials[1–7].

| Materials | $\rho$ (kg/m$^3$) | $k$ (W/m.K) | $C$ (J/kg.K) | $c$ (m/s) | $\alpha$ (dB/cm/MHz) | $\omega_b$ (kg/m$^3$s) |
|---|---|---|---|---|---|---|
| Tumor | 1050 | 0.54 | 3852 | 1509 | 0.57 | 0.515 |
| Skin | 1000 | 0.3425 | 3155 | 1541.6 | 1.6694 | --- |
| Water | 998 | 0.598 | 4184 | 1484 | 0.0021 | --- |
| Blood | 1050 | --- | 3617 | --- | --- | --- |

Supplemental Table 3. Driving parameters, hydrophone calibration in water, and simulated focal pressure within tumor tissue.

| Case | Frequency (MHz) | Input pressure amplitude (MPa) | Hydrophone focal P-P (water) (MPa) | Simulated focal P-P (tumor) (MPa) | Sonication time (s) |
|---|---|---|---|---|---|
| Low | 3.78 | 0.02624 | 4.2815 | 3.8393 | 15 |
| Medium | 3.78 | 0.03742 | 6.1037 | 5.4751 | 15 |
| High | 3.78 | 0.04645 | 7.5778 | 6.7963 | 15 |

**Supplemental Figure 1: Overview in silico 3D computational model rendering and geometry.** A) Schematic of the simulation geometry. B) Grid independence study based on peak-to-peak pressure at the acoustic focus.

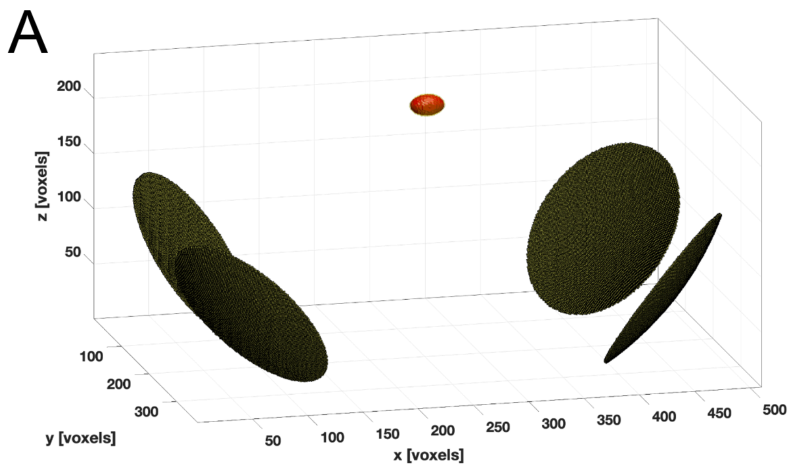

**Supplemental Figure 2: *In silico* simulation of T-FUS$_{Mid}$ treatment showing an increase in temperature and ablation volume as sonication period increases.** A-B) 3D computational model rendering temperature distributions through tumor center in xz, yz, and xy planes after T-FUS$_{Mid}$ treatment, with sonication period of 5 s and 25 s, respectively. C-D) 3D thermal dose (CEM43) maps through tumor center in xz, yz, and xy planes after T-FUS$_{Mid}$ treatment, with sonication period of 5 s and 25 s, respectively.

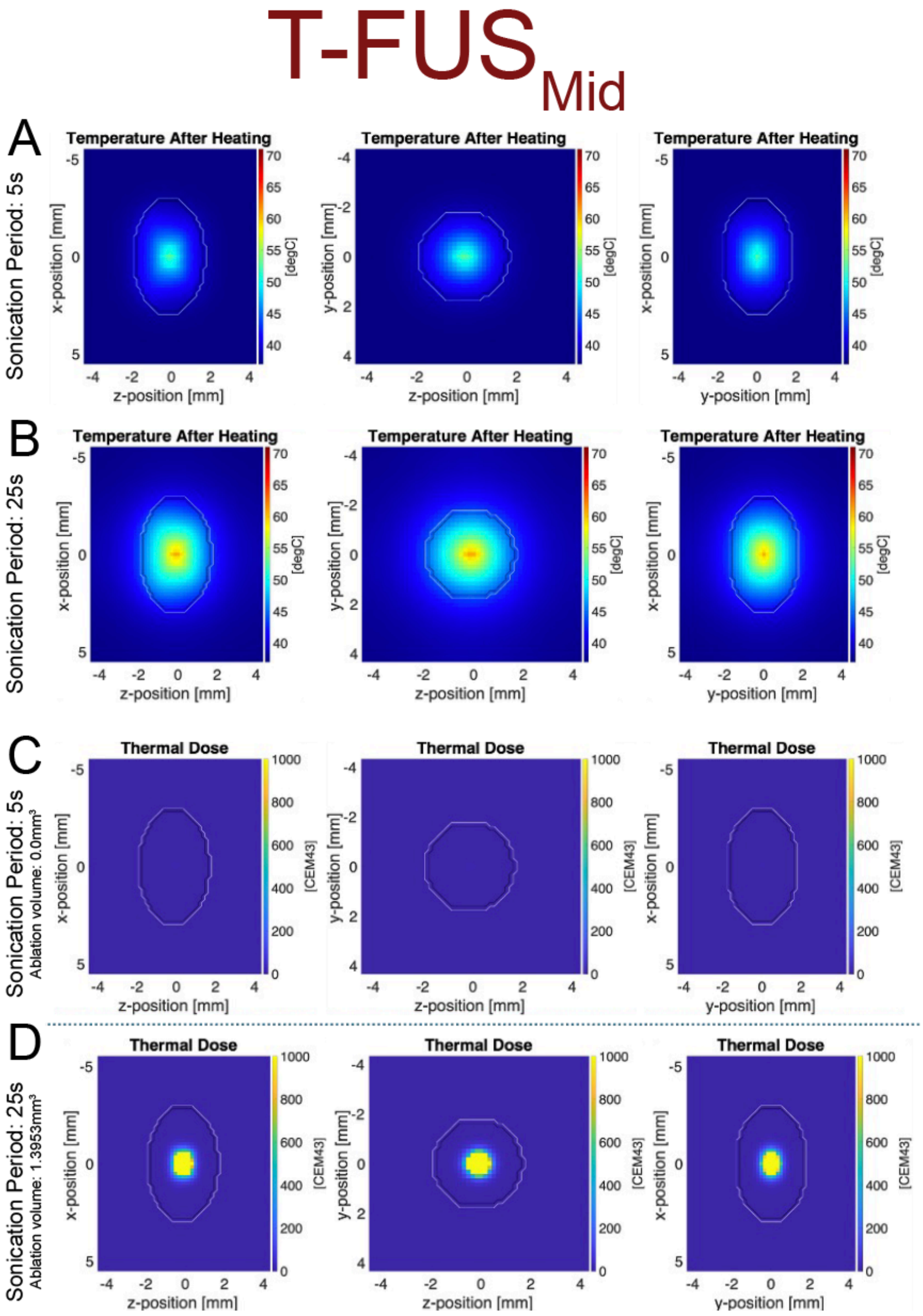

**Supplemental Figure 3: T-FUS ablation does not alter peripheral biodistribution of $[^{89}Zr]$-αCD47 in 4T1 tumor-bearing mice.** A-F) $[^{89}Zr]$-αCD47 uptake in blood, heart, spleen, kidney, lung, and tumor draining lymph node (TDLN) three days following sham or T-FUS treatment. G) Tissue-to-blood ratio of $[^{89}Zr]$-αCD47 uptake in peripheral organs at endpoint. Significance assessed by unpaired t-test. n.s. = not significant.

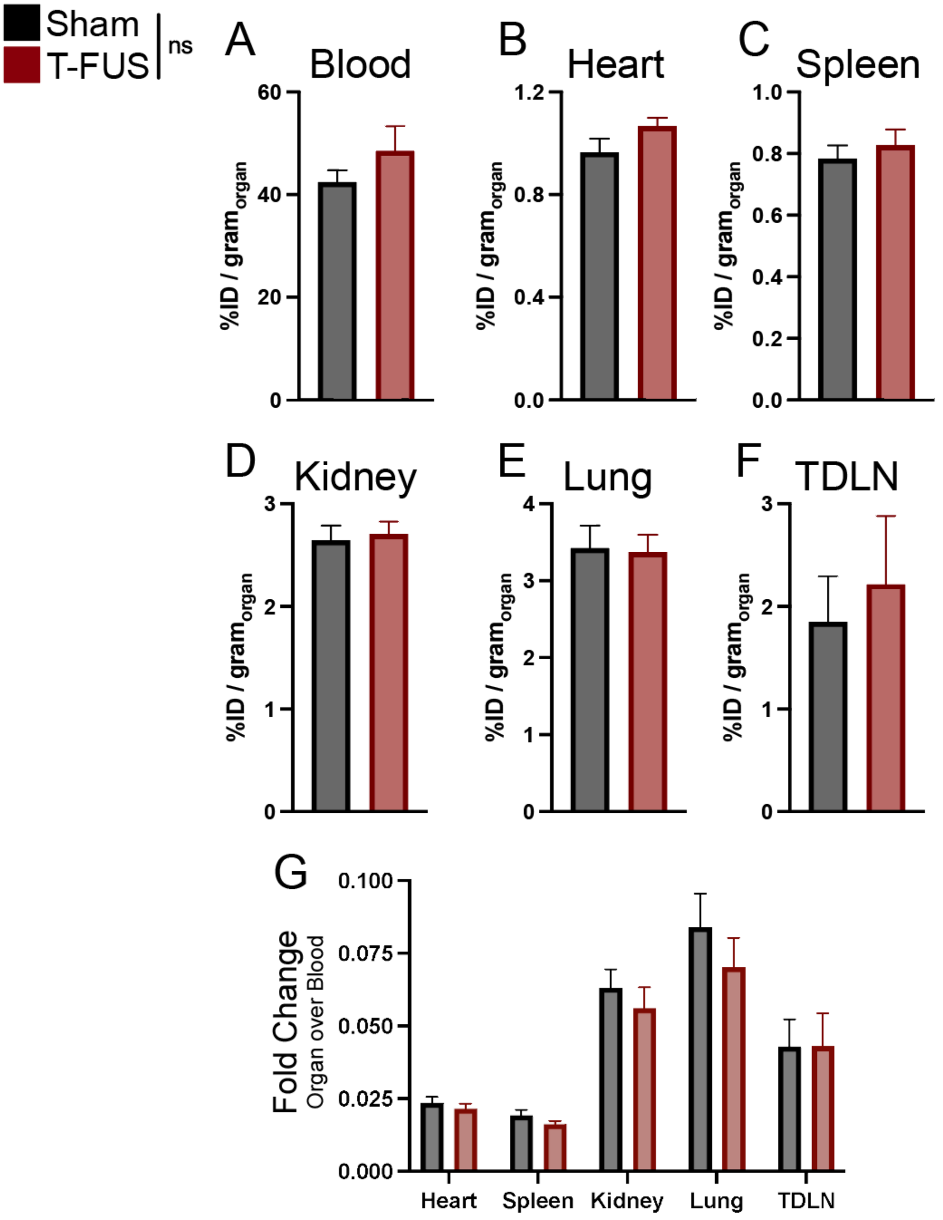

**References:**


1. Rahpeima, R. & Lin, C.-A. A comprehensive numerical procedure for high-intensity focused ultrasound ablation of breast tumour on an anatomically realistic breast phantom. *PLoS ONE* **19**, e0310899 (2024).
2. Martin, E., Jaros, J. & Treeby, B. E. Experimental Validation of k-Wave: Nonlinear Wave Propagation in Layered, Absorbing Fluid Media. *IEEE Trans. Ultrason., Ferroelect., Freq. Contr.* **67**, 81–91 (2020).
3. Bottiglieri, A., Sheth, R. A. & Prakash, P. A Computational Modeling Approach to Investigate the Influence of Hyperthermia on the Tumor Microenvironment. *JoVE* 65870 (2023) doi:10.3791/65870.
4. J.R. Speakman. Obesity and thermoregulation. in *Handbook of Clinical Neurology* vol. 156 431–443 (Elsevier, 2018).
5. Faber, P. & Garby, L. Fat content affects heat capacity: a study in mice. *Acta Physiologica Scandinavica* **153**, 185–187 (1995).
6. Jaunich, M., Raje, S., Kim, K., Mitra, K. & Guo, Z. Bio-heat transfer analysis during short pulse laser irradiation of tissues. *International Journal of Heat and Mass Transfer* **51**, 5511–5521 (2008).
7. Bhagat, P. K., Kerrick, W. & Ware, R. W. Ultrasonic characterization of aging in skin tissue. *Ultrasound in Medicine & Biology* **6**, 369–375 (1980).
